# Convergent Sequence Features of Antiviral B Cells

**DOI:** 10.1101/2023.09.06.556442

**Authors:** Alexandra A. Abu-Shmais, Matthew J. Vukovich, Perry T. Wasdin, Yukthi P. Suresh, Scott A. Rush, Rebecca A. Gillespie, Rajeshwer S. Sankhala, Misook Choe, M. Gordon Joyce, Masaru Kanekiyo, Jason S. McLellan, Ivelin S. Georgiev

**Affiliations:** Vanderbilt Vaccine Center, Vanderbilt University Medical Center, Nashville, TN 37232, USA; Department of Pathology, Microbiology and Immunology, Vanderbilt University Medical Center, Nashville, TN 37232, USA; Program in Chemical and Physical Biology, Vanderbilt University Medical Center; Nashville, TN 37232, USA; Department of Molecular Biosciences, The University of Texas at Austin, Austin, TX 78712, USA; Vaccine Research Center, National Institute of Allergy and Infectious Diseases, National Institutes of Health, Bethesda, MD 20892, USA; Emerging Infectious Disease Branch, Walter Reed Army Institute of Research, Silver Spring, MD 20910, USA; Henry M. Jackson Foundation for the Advancement of Military Medicine, Inc., Bethesda, MD 20910, USA; Vanderbilt Institute for Infection, Immunology and Inflammation, Vanderbilt University Medical Center, Nashville, TN 37232, USA; Department of Computer Science, Vanderbilt University, Nashville, TN 37232, USA; Center for Structural Biology, Vanderbilt University, Nashville, TN 37232, USA; Program in Computational Microbiology and Immunology, Vanderbilt University Medical Center; Nashville, TN, 37232, USA

## Abstract

Throughout life, humans experience repeated exposure to viral antigens through infection and vaccination, building diverse antigen-specific antibody repertoires. In recent years, these repertoires have become an important source for novel antibody-based antiviral therapeutics, yet there is still limited understanding of the determinants of antibody-antigen specificity. Here, we generated a large dataset mapping antibody sequence to antigen specificity for thousands of B cells, by screening the repertoires of a set of healthy individuals against twenty viral antigens representing diverse pathogens of biomedical significance. Analysis revealed antigen-specific patterns in variable gene usage, gene pairing, and somatic hypermutation, as well as the presence of convergent antiviral signatures across multiple individuals. These results help define the characteristics of human antibody repertoires simultaneously against an unprecedented number and diversity of viral targets. Understanding the fundamental rules of antibody-antigen interactions can lead to transformative new approaches for the development of antibody therapeutics and vaccines against current and emerging viruses.

## INTRODUCTION

The B cell compartment of the adaptive immune system mediates a critical role in the generation of antibodies against invading pathogens [1]. A principal feature of antibodies and their membrane-bound counterpart, the B cell receptor (BCR), is the exquisite specificity displayed against cognate antigens. Indeed, the human antibody repertoire is subject to exceptional levels of diversity [2, 3], acting as an indispensable protective mechanism against the myriad of infectious diseases we may encounter throughout life. Yet still, the phenomenon of public antibody clonotypes – highly similar antibodies identified in multiple individuals – has been described in several settings [4–7], challenging this dogma of virtually unlimited antibody diversity. Uncovering convergent antibody signatures present in antiviral repertoires can help elucidate canonical motifs of common immune responses and can guide vaccine design aiming to elicit a desired antibody clonotype at the population level.

Importantly, antigen specificity and therefore diversity of the antibody repertoire is the product of several complex and spatiotemporal mechanisms [3]. Combinatorial and junctional diversity estimates of the naïve repertoire alone are upwards of 10^12^ BCR sequences and increase in orders of magnitude when considering the memory B cell compartment [2, 8, 9]. Upon antigen encounter, B cells travel to distal germinal centers and undergo cyclic rounds of somatic hypermutation resulting in the generation of highly specific antibodies that are able to recognize new and repeated pathogenic attacks. Generally, this is an idiosyncratic process, however, in the setting of a public antibody clonotype, this mechanism is occurring in a highly convergent manner.

BCR specificity can be defined by several sequence metrics, owing to the complex mechanisms described. Namely: germline variable (V), diversity (D), and joining (J) gene use, the length and amino acid composition of the complementarity determining region 3 (CDR3), the degree of somatic hypermutation across the heavy and light chain, the constant region isotype, and clonality. The precise definition of a public antibody clonotype can vary, with different restrictions on sequence identity and/or heavy and light chain gene usage. A prevalent definition of “public” requires that clonotypes be present in more than one individual, sharing the same variable heavy (VH) genes and displaying highly similar CDRH3 regions, although with the advance of paired heavy-light chain sequencing, analogous restrictions on the variable light (VL) genes and CDRL3 sequences can also be added. Convergent antibody signatures have been reported in several anti-pathogen repertoires, including SARS-CoV-2 [10–13], human immunodeficiency virus 1 (HIV-1) [14], dengue [15, 16], respiratory syncytial virus (RSV) [17], ebola [18, 19], influenza [20–23], and others. The selective pressure towards expansion of a public clonotype suggests certain germline gene segments may have inherent features that facilitate recognition of these viral proteins. Along this line, convergent sequence features identified within a specific antiviral repertoire, irrespective of public clonality, can lend further support to the notion of canonical molecular and structural features mediating recognition of antigenic targets.

Advances in repertoire sequencing [24–28] have enabled a greater understanding of the relationship between BCR sequence and antigen specificity, in the context of viral infectious disease. Particularly, since the onset of the COVID-19 pandemic, several studies have shed light on SARS-CoV-2 specific BCR signatures [10–13, 29–31]. However, the vast majority of published BCR sequences in the public domain lack information about their cognate antigens, as even high-throughput antibody sequence identification methods, such as next-generation sequencing, are generally decoupled from the process of antibody functional characterization. In default of large datasets of antigen annotated BCR sequences representing multiple viral antigen specificities, the similarities and differences of antibody responses to different viral pathogens remains unclear.

In this study, leveraging a high-throughput B cell discovery technology termed LIBRA-seq, Linking B cell Receptor to Antigen Specificity, we systematically analyzed antigen specific B cell sequences against twenty viral antigens, representing several pathogens of current biomedical interest. Patterns in antibody chain usage, variable gene usage, CDRH3 length, and somatic hypermutation frequency were observed with respect to viral antigen specificity. Such patterns are evident when restricting analysis to B cell breadth, as opposed to specificity, suggesting intrinsic differences between mono-reactive and cross-reactive B cells exist. Moreover, analysis revealed the presence of hundreds of antiviral public clonotypes, clustered on the basis of matching CDR3 length, a minimum of 70% amino acid identity in the CDR3 region, and matching variable and joining genes in both the heavy and light chains. These data demonstrate that convergent B cell molecular features can be readily identified across multiple individuals and additionally are occurring at a public antibody level.

## RESULTS

### Donor peripheral blood mononuclear cells (PBMCS) and LIBRA-seq viral antigen screening library

Donor PBMC samples were collected between June 2021 and May 2022. An equal distribution of men and women ages 23 through 59 were used in this study (Supplementary table 1). Participants are tested for HIV-1/-2 and hepatitis B/C within 90 days of collection date and are assumed to be healthy, lacking indication of infectious morbidities, at the time of sample collection.

LIBRA-seq is a high throughput antibody discovery tool that enables the simultaneous detection of B cell receptor sequence and antigen specificity from a single biological sample. Here, the antigen screening library included twenty viral antigens associated with common infections and vaccinations, as well as a small subset of other antigens of current biomedical significance (Figure 1). The library incorporated major antigenic targets for the selected pathogens, and in some cases multiple strains per pathogen were included. Antigens have been grouped by viral family: Coronaviridae, Paramyxoviridae, Pneumoviridae, Orthomyxoviridae, and Flaviviridae. Sorted B cells displaying binding to HIV-1 BG505 Env, a negative control antigen that is not expected to be recognized in healthy individuals, were removed from all datasets on the basis of cell promiscuity.

**Figure 1:**
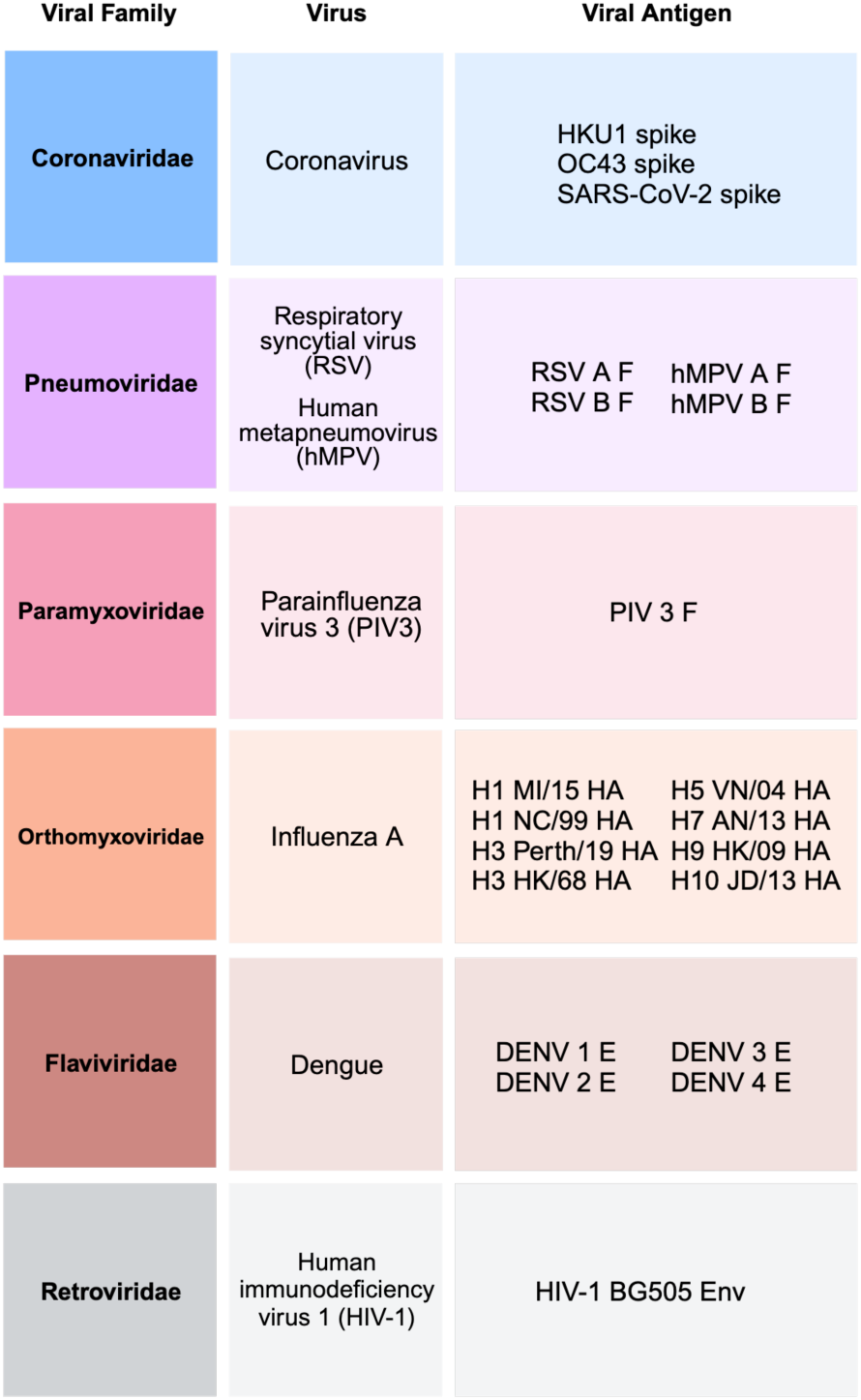
LIBRA-seq viral antigen screening library. The screening library included antigens associated with common infections and vaccinations, as well as a selection of other antigens of biomedical significance. A total of twenty antigens representing multiple strains across five distinct viral families was included. This antigen library incorporates some of the major antigenic targets for the selected pathogens. Viral families are shown in darker shaded boxes, followed by virus and viral antigen in lighter shaded boxes. HIV-1 BG505 env served as negative control antigen and was not used for system wide analyses.

### Generation of antigen-specific B cell datasets by single-cell sequencing, analytics, and recombinant validation

We sorted CD19+, IgG+, antigen +, B cells from ten healthy donor PBMC samples using fluorescently tagged LIBRA-seq antigens as probes (Figure 2A). Resulting paired heavy and light chain sequences, along with LIBRA-seq scores which provide a relative measure of antigen binding, were analyzed as described in the methods. Antigen specificity was assigned based on the LIBRA-seq predictions, where a cell that displayed a LIBRA-seq score of one or greater for antigen X, is considered to bind antigen X. After filtering the data for sequences that displayed promiscuous binding patterns, i.e binding to HIV-1 BG505 or binding to multiple antigens belonging to more than one viral family, we obtained a total of 2711 B cell receptor (BCR) sequences with a LIBRA-seq score of at least one for a minimum of one antigen. Of those sequences, 2175 were IgG restricted, dominated by the IgG1 subclass. Coronaviridae represented the largest group of antigen-specific B cells, followed by Pneumoviridae, Orthomyxoviridae, Paramyxoviridae, and Flaviviridae (Figure 2B). The population of IgA and IgM recovered cells can be attributed to events that fall near the IgG+/FITC+ demarcation gate during flow cytometric sorting.

**Figure 2:**
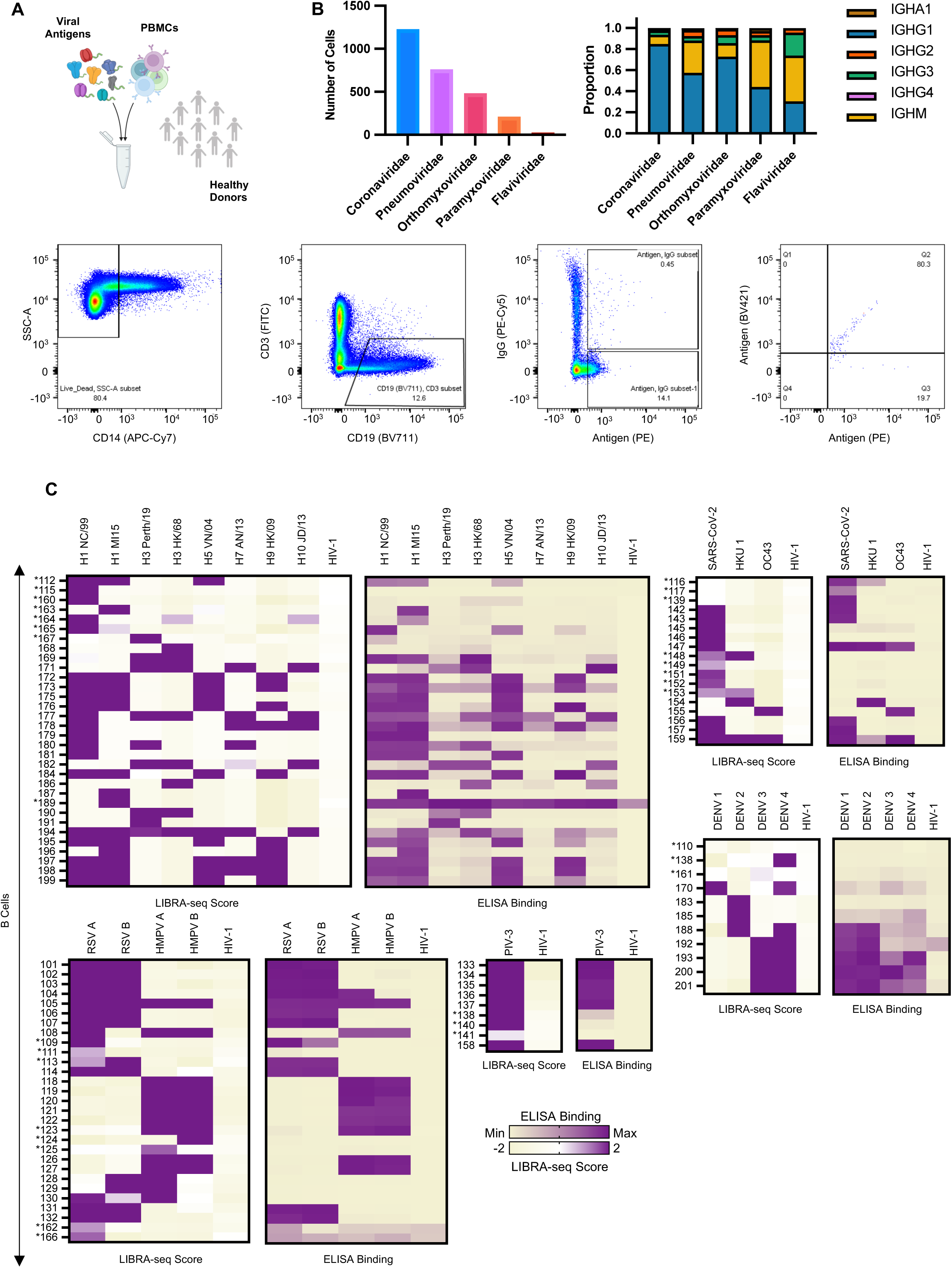
Identification of antigen-specific B cells from healthy donors. 2A: Schematic of sample processing and flow cytometric staining to identify antigen specific B cells. Fluorescence-activated cell sorting of lymphocytes, B cells, and antigen specific IgG cells. Cells were stained with anti-CD14 antibody conjugated to allophycocyaninin cyanine dye (APC-Cy7), anti-CD19 conjugated to brilliant violet 711 (BV711), anti-CD3 conjugated to fluorescein isothiocyanate (FITC), anti-IgG conjugated to phycoerythrin cyanine dye (PE-Cy5). LIBRA-seq antigens were biotinylated and conjugated to streptavidin-phycoerythrin (Strep-PE) and streptavidin brilliant violet 421 (BV421). 2B: Number of B cells identified for each antigen specificity category after computational filtering. Proportion of immunoglobulin isotypes identified within each antigen specificity category. 2C: LIBRA-seq scores (left) and ELISA binding (right) of recombinantly expressed monoclonal antibodies, partitioned by viral family. Antibodies are numbered 100-201. Antibodies denoted with asterisk originate from the unfiltered dataset. Some antibodies with asterisk are listed twice if binding predictions span viral families. LIBRA-seq scores for each antigen are displayed as a heatmap with a LIBRA-seq score of −2 displayed as beige, 0 as white, and a LIBRA-seq score of 2 as purple; scores lower or higher than that range are shown as −2 and 2, respectively. ELISA binding, calculated as absorbance at 450 nm, is displayed as a heatmap with the minimum detected signal (0.03) displayed as beige and maximum detected signal (3.5) displayed in purple.

To provide a measure of the robustness of the LIBRA-seq antigen specificity predictions, we selected 69 paired heavy and light chain sequences from our filtered dataset to express as monoclonal antibodies. B cells were prioritized based on binding profile and LIBRA-seq score. Additionally, we selected 30 B cells from the subset of sequences that did not meet our filtering criteria (i.e., B cells with promiscuous binding patterns), for a total of 99 antibodies for recombinant expression and validation. Of the 69 antibodies ordered from the filtered datasets, 59 of them bound to at least one of the antigens it was predicted to bind by LIBRA-seq. From the subset of sequences classified as promiscuous, 10 out of the 30 bound as predicted without binding to HIV-1 BG505 and were therefore moved to the filtered dataset (Figure 2C).

### Antigen-specific B cell patterns in V gene usage, IGHV:IGL(K)V gene pairings, CDRH3 length, and somatic hypermutation

We next explored the immunoglobulin heavy and light chain sequence features of the IgG restricted sequences and excluded Flaviviridae reactive B cells due to insufficient numbers. Analysis revealed usage of 47 heavy chain, IGHV, and 56 light chain, IGL(K)V, variable genes, across the dataset with several IGHV:IGL(K)V gene patterns occurring at a higher frequency within certain antigenic categories. For example, IGHV5-51:IGLV1-40 and IGHV5-51:IGLV2-23 were frequently observed within Paramyxoviridae and did not overlap with other specificity categories (Figure 3A-B). In accordance with germline complexity of heavy chain variable segment families and published findings on VH bias of productive peripheral B cells [32, 33]; all of the antigenic specificity categories exhibited a preference towards the VH3 family, while VH6 and VH7 were exceedingly rare. Notably, 21.2% of Paramyxoviridae-specific cells belonged to the VH5 family, an overrepresentation in comparison to the other categories. A consistent bias across all categories towards IGHJ4 was also observed (Supplementary Figure 3). IGHV1-69 has been implicated in several anti-pathogen repertoires [34–40] and indeed was observed among all specificity categories here. In comparing to previously published repertoires, unselected by any antigen specificity [41, 42], several genes were found at a higher frequency in the antigen specificity categories, most notably IGHV1-18 and IGHV3-30. (Figure 3A). A similar distribution of heavy chain complementarity determining region 3 (CDRH3) lengths among the specificity categories was observed with Coronaviridae, Pneumoviridae, Orthomyxoviridae, and Paramyxoviridae having a majority of 16, 16, 18, and 12 amino acid long CDRH3 regions, respectively (Table 1, Supplementary Figure 1A). Finally, we evaluated the proportion of heavy chain variable (VH) somatic hypermutation (1-VH identity calculated at the amino acid level) and found that the Pneumoviridae category displayed the highest frequency of VH somatic hypermutation with an average VH identity of 0.93 and a range of 0.84 -1.00 (Figure 3C, Table 1). Interestingly, VH somatic hypermutation correlated with VL somatic hypermutation for all antigen specificity categories; Coronaviridae (p= 7.2^-63^, Spearman 0.47), Pneumoviridae (7.9^-48^, Spearman 0.57), Paramyxoviridae (p=2.1^-21^, Spearman 0.57), Orthomyxoviridae (p= 8.7^-47^, Spearman 0.62) (Supplementary Figure 1C).

**Figure 3:**
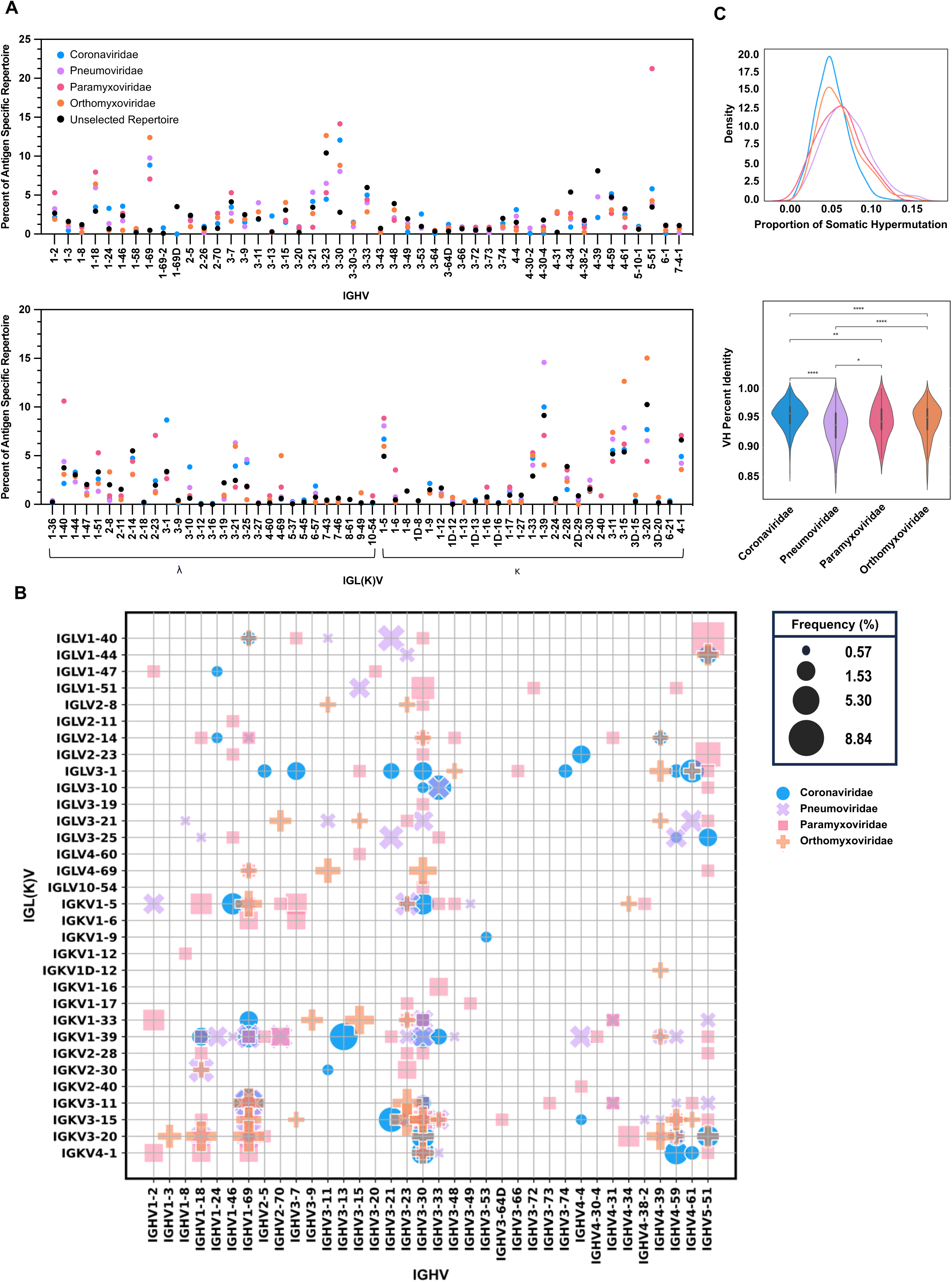
Diversity of viral antigen specific B cells. 3A: IGHV and IGL(K)V gene usage in B cells specific to Coronaviridae (blue), Pneumoviridae (purple), Paramyxoviridae (pink), and Orthomyxoviridae (orange) antigens. IGHV gene usage from previously published, unselected repertoires shown in black. Gene usage is represented as frequency of gene use within antigen specificity category. Circles are superimposed. Zeros are not plotted. 3B: The frequency of different IGHV:IGL(K)V gene pairs for B cells specific to Coronaviridae (blue circle), Pneumoviridae (purple cross), Paramyxoviridae (pink square), and Orthomyxoviridae (orange plus) antigens. The size of each data point represents the frequency of the corresponding IGHV:IGK(L)V pair within its antigen specificity category. Frequencies above 0.5% are represented. 3C: Frequency of heavy chain variable (VH) somatic hypermutation represented as 1-VH identity calculated at the amino acid level. Violin plot width is proportional to the fraction of B cells with the indicated proportion of VH somatic hypermutations. Two sided Mann-Whitney-Wilcoxon test with Bonferroni correction used to calculate significance. Coronaviridae vs. Pneumoviridae: p = 5.626^-34^, Pneumoviridae vs. Paramyxoviridae: p = 3.990^-02^, Paramyxoviridae vs. Orthomyxoviridae:p = 1.000 Coronaviridae vs. Paramyxoviridae: p = 1.903^-03^, Pneumoviridae vs. Orthomyxoviridae: p = 2.173^-06^, Coronaviridae vs. Orthomyxoviridae: p = 2.751^-07^.

**Table 1:** Sequence metrics of IgG restricted and public clonotype datasets. Total cell counts, mean/minimum/maximum/mode heavy chain complementary determining region 3 (CDRH3) lengths, and mean/minimum/maximum/mode variable heavy (VH) somatic hypermutation (SHM) values for IgG restricted, public clonotype, and reactivity pattern restricted datasets.

| IgG Restricted | Coronaviridae | Pneumoviridae | Paramyxoviridae | Orthomyxoviridae |  |
| --- | --- | --- | --- | --- | --- |
| Total cells | 1119 | 521 | 113 | 419 |  |
| Mean CDRH3 length | 17 | 17 | 16 | 18 |  |
| Minimum CDRH3 length | 7 | 7 | 8 | 8 |  |
| Maximum CDRH3 length | 37 | 29 | 27 | 30 |  |
| Mode CDRH3 length | 16 | 16 | 12 | 18 |  |
| Mean VH Percent Identity | 0.95 | 0.93 | 0.94 | 0.94 |  |
| Minimum VH Percent Identity | 0.85 | 0.84 | 0.87 | 0.84 |  |
| Maximum VH Percent Identity | 1.00 | 1.00 | 1.00 | 0.99 |  |
| Mode VH Percent Identity | 0.95 | 0.94 | 0.94 | 0.96 |  |
| Public Clonotypes | Coronaviridae | Pneumoviridae | Paramyxoviridae | Orthomyxoviridae | Flaviviridae |
| Total cells | 358 | 67 | 27 | 32 | 16 |
| Mean CDRH3 length | 11 | 16 | 17 | 15 | 17 |
| Minimum CDRH3 length | 6 | 12 | 15 | 10 | 15 |
| Maximum CDRH3 length | 23 | 24 | 20 | 21 | 18 |
| Mode CDRH3 length | 6 | 16 | 19 | 14 | 18 |
| IgG Restricted by Reactivity Pattern | Mono-reactive | Cross-reactive |  |  |  |
| Total cells | 1713 | 459 |  |  |  |
| Mean CDRH3 length | 17 | 17 |  |  |  |
| Minimum CDRH3 length | 7 | 8 |  |  |  |
| Maximum CDRH3 length | 37 | 30 |  |  |  |
| Mode CDRH3 length | 16 | 18 |  |  |  |
| Mean VH Percent Identity | 0.95 | 0.93 |  |  |  |
| Minimum VH Percent Identity | 0.84 | 0.84 |  |  |  |
| Maximum VH Percent Identity | 1.00 | 0.99 |  |  |  |
| Mode VH Percent Identity | 0.95 | 0.92 |  |  |  |

### Identification of public viral antigen-specific B cells

To interrogate the existence of public antibodies within our dataset, we built a reference library of paired BCR sequences from published sources [41, 42]. The reference library contained 1,060,594 human sequences categorized into five donor bins. As CoVAbDab is a consolidation of published and patented antibody sequences against beta coronaviruses and donor identity was not uniformly assigned at the time of data generation, the 12,021 human BCR sequences from the database used towards our reference library were categorized into a single bin. While this represents an oversimplification, it eliminates the possibility of inflated donor assignment. To increase the likelihood of identifying public clonotypes and explore the propensity of IgA and IgM antibodies towards the induction of population level antiviral responses, all sequences from our dataset were used in comparison against the reference library. Sequences were screened for matching CDRH3 and CDRL3 amino acid length, 70% minimum amino acid identify in both CDR3 regions, and matching variable and joining gene use in both heavy and light chains. From the LIBRA-seq datasets, 113 sequences matched with 387 sequences in the reference library (Supplementary Table 2), meeting the similarity criteria described above; these sequence matches were used for all subsequent public antibody analysis.

### Public antibody clonotypes of different viral antigen specificities have distinct V gene usage and IGHV:IGL(K)V gene patterns

Analysis of the IGHV and IGL(K)V gene use among viral antigen public clonotypes revealed localization across 30 variable heavy genes and 24 variable light genes. Despite the isolation of low cell numbers for Flaviviridae reactive cells, 16 public clonotypes were identified through our similarity criteria, and were therefore included in subsequent analysis. Interestingly, the most frequently observed heavy chain variable genes within Coronaviridae IGHV 4-59 (48.04%), Pneumoviridae IGHV3-33 (22.3%), Paramyxoviridae IGHV3-53 (18.51%), and Orthomyxoviridae IGHV3-33 (31.2%), differed from the most frequently leveraged genes by each category within the IgG restricted dataset (Figure 3A, Figure 4A). Of the Coronaviridae public clonotypes identified, several of the heavy chain genes have previously been reported in SARS-CoV-2 public B cells: IGHV3-30, [12], IGHV3-53, IGHV1-58, (Wang et al., 2022), IGHV3-66 [30], IGHV1-69, IGHV4-59, and IGHV3-7 [10]. Other previously identified public heavy chain genes in various anti-pathogen repertoires were also observed: IGVH 1-18 [21] and IGVH 1-69 [36] among influenza A hemagglutinin (HA) binding B cells, and IGVH 1-18, IGVH2-70, IGVH3-21, IGVH3-9, and IGVH 1-2 (Mukhamedova et al., 2021) within RSV repertoires. As revealed with IGHV gene analysis, the most frequently observed light chain variable genes within Coronaviridae IGKV 3-20 (52.79%), Pneumoviridae IGKV2-30 (16.4%), Paramyxoviridae IGKV 2-28 (25.9%), and Orthomyxoviridae IGLV3-10 (21.8%) also differed from the most frequently observed genes in the IgG restricted dataset. Moreover, a smaller distribution of IGL(K)V genes, 24 opposed to 56, are leveraged across the viral antigen public clonotypes, suggesting a role for light chain coherence-a phenomenon recently described [41]. Analysis of IGHV:IGL(K)V gene pairing frequencies across the antigenic specificities revealed previously known gene usage patterns, such as IGHV 4-59: IGKV3-20 for SARS-CoV-2 [11] within Coronaviridae and IGHV 1-18: IGKV2-30 for RSV A/B [17] within Pneumoviridae, as well as enrichments for, to our knowledge, not yet reported, gene patterns and antigen specificities among public clonotypes, such as IGHV5-51: IGKV4-1, IGHV3-53: IGKV3-20, and IGHV1-18: IGKV2-28 for PIV-3 within Paramyxoviridae, and IGHV 3-7:IGKV 1-6 for DENV 4 within Flaviviridae (Figure 4B, Supplementary Table 3).

**Figure 4:**
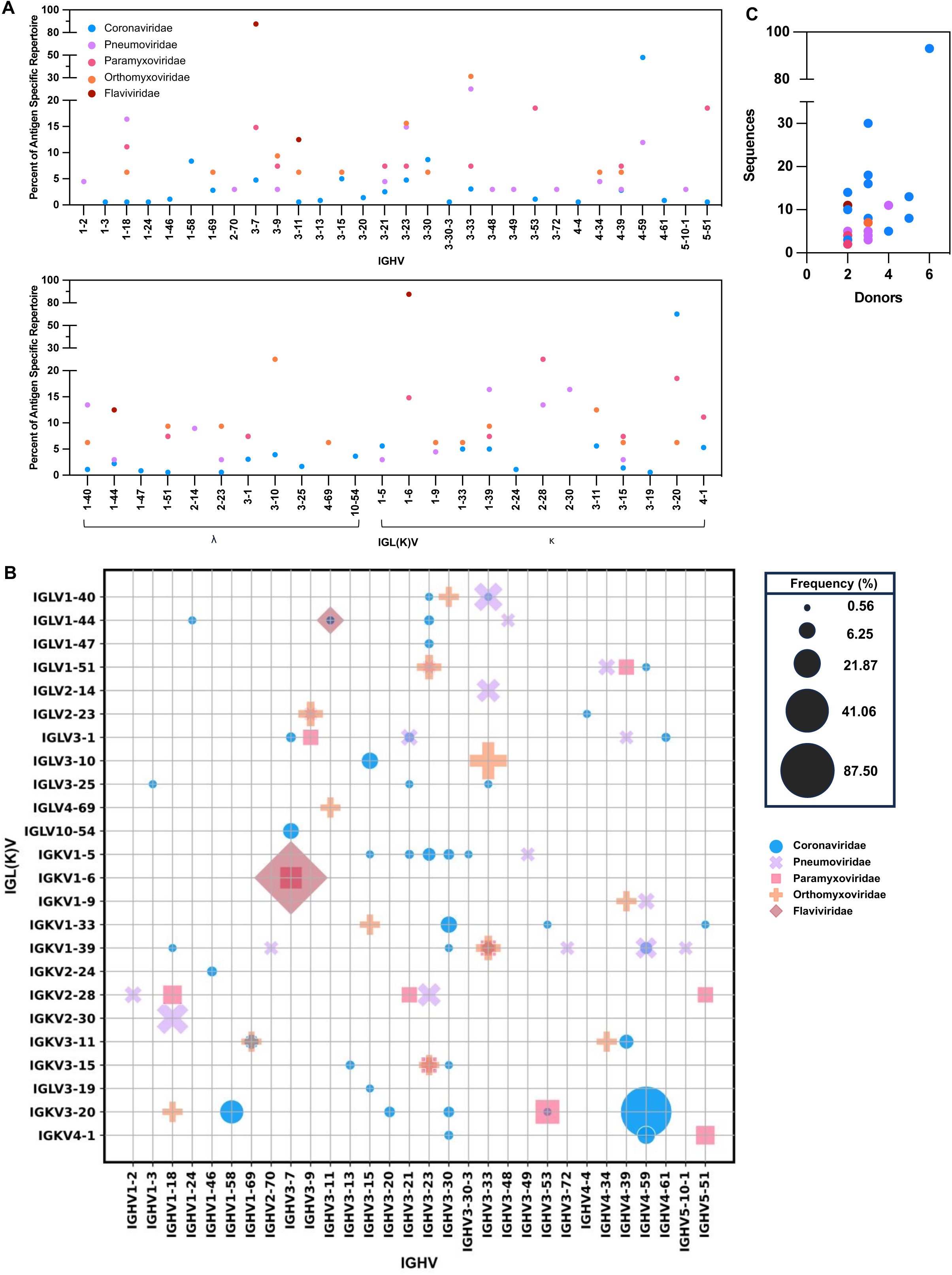
B cell characteristics of viral antigen public clonotypes. 4A: IGHV and IGL(K)V gene usage in public clonotypes specific to Coronaviridae (blue), Pneumoviridae (purple), Paramyxoviridae (pink), Orthomyxoviridae (orange), and Flaviviridae (red) antigens. Gene usage is represented as frequency of gene use within antigen specificity category. Circles are superimposed. Zeros are not plotted. 4B: The frequency of different IGHV:IGL(K)V gene pairs for public clonotypes specific to Coronaviridae (blue circle), Pneumoviridae (purple cross), Paramyxoviridae (pink square), Orthomyxoviridae (orange plus), Flaviviridae (red diamond) antigens. The size of each data point represents the frequency of the corresponding IGHV:IGK(L)V pair within its antigen specificity category. Frequencies above 0.5% are represented. 4C: Clusters of public clonotypes within each antigen specificity category identified on similarity criteria of matching CDRH3 and CDRL3 amino acid length, 70% minimum amino acid identify in both CDR3 regions, and matching variable and joining gene use in both heavy and light chains. Y axis represents the total number of sequences within each cluster. X axis represents the total number of individuals each cluster was identified within. Colored as in previous (4A).

Expectedly, broadening the similarity margin to advise on heavy chain elements alone led to the identification of thousands of public clonotypes (Supplementary Table 2). The number of different IGHV and IGL(K)V germline genes leveraged by these public clones likewise increased, resulting in several IGHV:IGL(K)V gene pairs that were not observed in the restrictive dataset (Supplementary Figure 4A-B). Collectively, these results demonstrate that public clonotypes of different antigen specificities leverage distinct patterns in variable gene usage and IGHV:IGL(K)V gene pairings.

### Convergent viral antigen-specific public clonotypes in healthy individuals

To investigate the presence of convergent molecular signatures in the public antibody response towards the antigens analyzed here, we performed a shared cluster analysis. Within the sequences identified as public, a shared cluster was defined as two or more sequences originating from a minimum of two donors that leverage the same pair of heavy/light variable and joining germline genes, while maintaining 70% amino acid identity and length in the CDR3 region. A total of 95 clusters were identified, of which 53, 18, 12, 10, and 2 were Coronaviridae, Pneumoviridae, Orthomyxoviridae, Paramyxoviridae, and Flaviviridae-specific, respectively (Figure 4C, Supplementary Table 3).

The largest cluster identified, cluster 81 IGHV4-59/HJ4: IGKV3-20/KJ1, was Coronaviridae-specific, and populated by 93 sequences across 6 donors. This cluster was characterized by an unusually short CDRH3 region (6 amino acids by IMGT numbering) and represents a previously identified public clonotype, by consideration of germline VH and VL gene use only, among S2 binding SARS-CoV-2-specific B cells [11]. Notably, cluster 80 IGH4-59/HJ2: IGKV3-20/KJ1 also represented a SARS-CoV-2-specific public clonotype and differed only in heavy chain joining germline gene usage, resulting in conservation of a leucine residue at the CDRH3 C terminus opposed to conservation of tyrosine at the same locus in cluster 81 (Figure 5). Analysis of shared clusters with a minimum of ten sequences revealed several other previously identified public clonotypes, such as cluster 3 IGHV1-18/HJ4: IGKV3-20/KJ2 among site V binding RSV-specific B cells following vaccination and natural infection [17, 40, 43], and clusters 9 IGHV1-58/HJ3: IGLV 3-20/KJ1[10] and 39 IGH3-30/HJ4: IGKV1-33/KJ3 [12] among SARS-CoV-2-specific B cells. Convergent molecular signatures in Dengue infection have been reported, however, analysis of IGHV:IGL(K)V gene usage patterns among public clonotypes have yet to be explored in detail. Here, cluster 61 IGHV3-7/HJ3:IGKV1-6/KJ1, specific to DENV 4, was identified in 3 unique donors and characterized by an 18 amino acid long CDRH3 region with less consensus between residues 7 through 12, potentially suggesting varied D gene use or preferential sites of somatic hypermutation. Among other Coronaviridae-specific shared clusters, cluster 62 IGHV3-7/J3:IGLV10-54/LJ3, OC43-specific, was identified in 5 donors and interestingly, represents the only public clonotype in this analysis leveraging IGLV10-54 (Figure 5, Supplementary Table 3).

**Figure 5:**
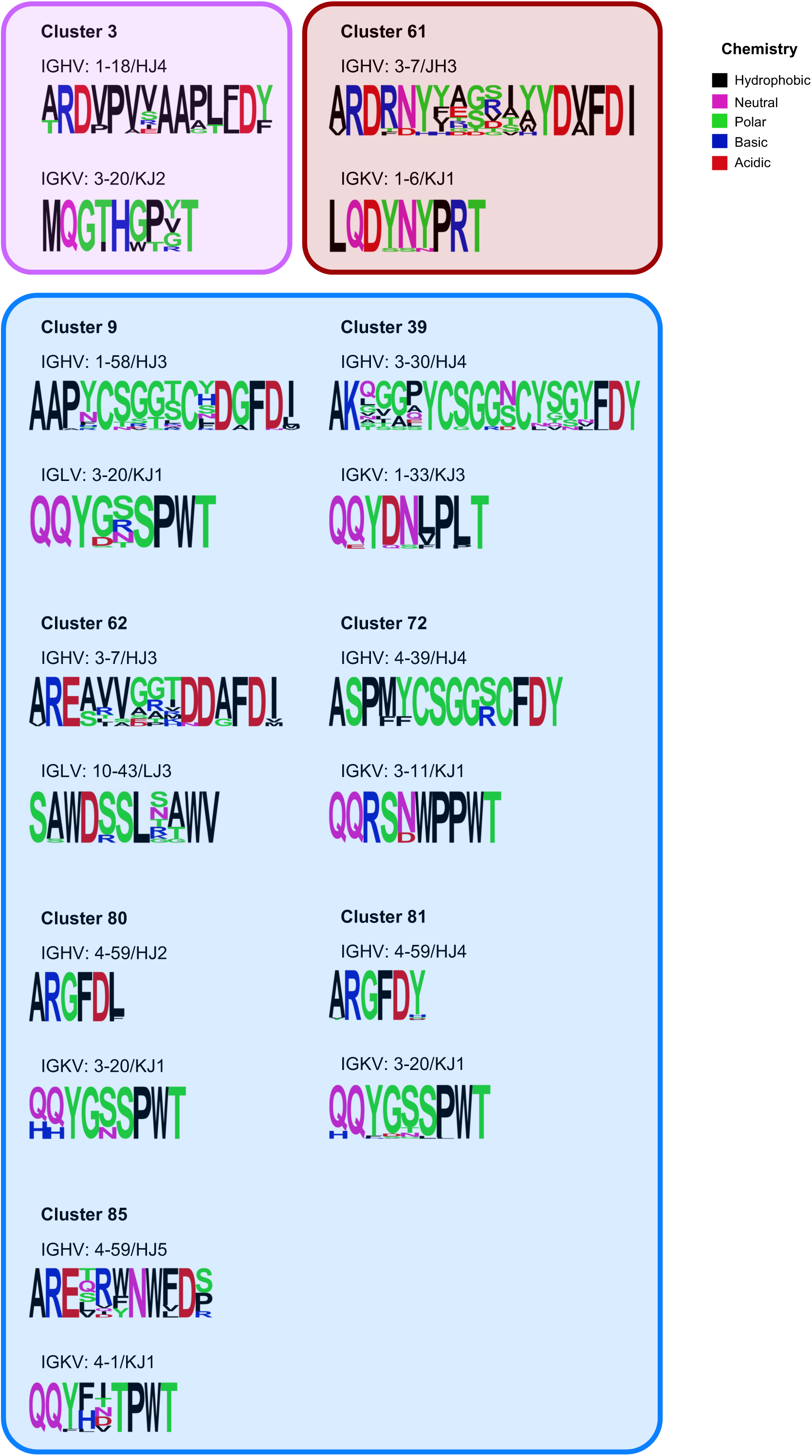
Antiviral public antibodies exhibit convergent CDR3 sequences. IGHV/HJ and IGL(K)V/L(K)J gene usage and CDRH3 and CDRL3 sequence are shown for clusters populated by a minimum of ten sequences across at least two donors. For each cluster, the CDRH3 and CDRL3 sequences are shown as a sequence logo, where the height of each letter represents the frequency of the corresponding amino-acid variant (single-letter amino acid code) at the indicated position. Clusters of the same antigen specificity are grouped in the same box. Blue indicates Coronaviridae, purple Pneumoviridae, and red Flaviviridae.

### Mono-reactive and cross-reactive B cells display distinct patterns in V gene usage and somatic hypermutation

Lastly, we investigated the relationship between the BCR sequence metrics observed above and B cell breadth. Here, a mono-reactive B cell was categorized as displaying a minimum LIBRA-seq score of one for a single antigen, while a cross-reactive B cell was defined as displaying a minimum LIBRA-seq score of one for at least two antigens within the screening library. From the IgG restricted dataset, 1713 cross-reactive and 459 mono-reactive sequences were identified (Table 1). While most IGHV and IGL(K)V genes were leveraged by both cell types, IGHV1-69 (16.1%) and IGHV3-30 (11.2%) were the most frequently observed IGHV genes for cross-reactive and mono-reactive cells, respectively, with IGKV1-39 (13.2% cross-, 8.8% mono-) observed most frequently for both categories (Figure 6A). Analysis of IGHV:IGL(K)V revealed several gene pairs specific to either cell type, such as IGHV1-69:IGKV3-20 among the mono-reactive subset and IGHV3-33:IGLV3-10 among the cross-reactive subset (Figure 6B). No statistical difference in the average amino acid length of the CDRH3 region was observed (data not shown), however, the majority of cross-reactive sequences displayed 18 amino acid long CDRH3 regions with a range between 8 and 30 residues while the majority of mono-reactive B cells displayed 16 amino acid long CDRH3 regions with a range between 7 and 37 residues (Table 1). Interestingly, our analysis revealed cross-reactive B cells displayed higher degrees of VH (p = 1.808^-36^) and VL (p = 3.353^-18^) somatic hypermutation than mono-reactive B cells (Figure 6C, Supplementary Figure 2A). As observed when analyzing somatic hypermutation with respect to antigen specificity in the IgG restricted dataset, VH somatic hypermutation moderately correlated with VL somatic hypermutation for both mono-reactive (p=1.4^-152^, Spearman 0.57) and cross-reactive cells (p=1.2^-36^, Spearman 0.54) (Supplementary Figure 2B). Analysis of VH percent identity across the most represented IGHV and IGLV genes observed in both cell types revealed higher degrees of somatic hypermutation in several IGHV genes among cross reactive cells despite being used less frequently than mono-reactive cells (Supplementary Figure 2C).

**Figure 6:**
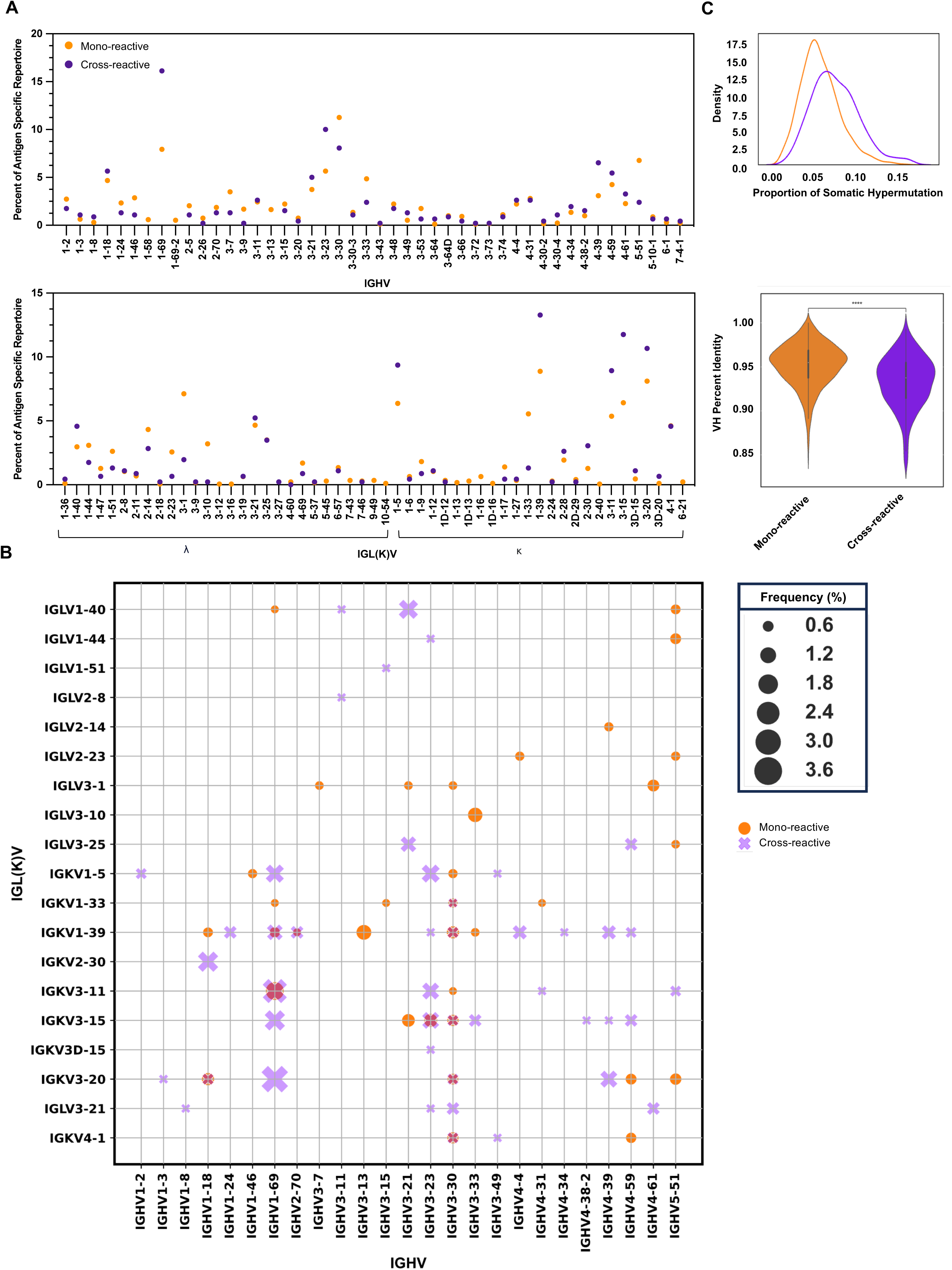
Molecular characteristics of mono-reactive and cross-reactive B cells. 6A: GHV and IGL(K)V gene usage in mono-reactive (orange) and cross-reactive (purple) B cells. Gene usage is represented as frequency of gene use within each reactivity category. Circles are superimposed. Zeros are not plotted. 6B:The frequency of different IGHV:IGL(K)V gene pairs for B cells of mono- and cross-reactivity. The size of each data point represents the frequency of the corresponding IGHV:IGK(L)V pair within its reactivity category. Frequencies above 0.5% are represented. 6C: Frequency of heavy chain variable (VH) somatic hypermutation represented as 1-VH identity calculated at the amino acid level. Violin plot width is proportional to the fraction of antibodies with the indicated proportion of VH somatic hypermutation. Kruskal-Wallis test with Bonferroni correction used to calculate significance. p = 1.808^-36^

## DISCUSSION

Leveraging LIBRA-seq, a high-throughput antibody discovery technology capable of screening a biological sample simultaneously against a theoretically unlimited number of targets, we performed a systematic analysis of antigen-specific B cell sequences against twenty viral antigens in healthy subjects. The inclusion of multiple pathogenic targets allowed us to explore patterns in several B cell receptor sequence metrics in the context of viral antigen reactivity, revealing differences in germline gene usage, heavy chain complementarity determining region 3 (CDRH3) length, and degree of somatic hypermutation. Through sequence analysis of the LIBRA-seq datasets, along with published antibody sequence data, we identified public antibody clonotypes against multiple targets, revealing convergent signatures in several antiviral pathogen repertoires. Together, our study represents the first of its kind to survey viral antigen-specific B cells at this scale, across multiple individuals, representing a crucial step forward in the generation of a comprehensive human antibody – viral antigen atlas.

The accepted definition of a public antibody clonotype varies in sequence identity and heavy and light chain gene usage. Owing to the contribution individual germline genes make towards the antigen binding interface, the induction of a public response can be driven solely by IGHV, as is the case for a subset of IGL(K)V – independent public antibodies [11, 44, 45]. Here, we imposed a stringent similarity criterion with respect to both heavy and light chain elements to increase confidence towards identification of recurring features contributing to the antiviral public response. As more antigen specificity data is obtained for public antibodies for various viral targets, the importance of including light chain vs. heavy chain only sequence features in defining antibody publicness will be further clarified.

Antigen specificity, indirectly afforded by BCR sequence, is the hallmark feature of antibodies. Identification of convergent motifs, both in public and private BCR sequences, lends support to the indication that germline gene segments may have inherent properties that enable recognition of viral proteins, such as the interactions between the aromatic residues within IGHV1-58 + IGHJ3 (public cluster 9) and SARS-CoV-2 RBD F486 [29] and the hydrophobic loop of IGHV1-69 and the hydrophobic pocket of influenza HA stem [46]. Unsurprising was the bias towards gene family VH3 within the IgG restricted dataset, as VH3 offers a repertoire of increased germline complexity not afforded by other VH segment families. Likewise, the preference towards IGHJ4 falls in line with previous findings and has been suggested to be the result of tyrosine desirability at the CDRH3 locus, by which IGHJ4 contributes two tyrosine residues [33, 47]. However, the increased usage of VH5 among Paramyxoviridae, in comparison to the other antigen specificity categories, was striking considering VH5 is comprised of two gene segments, suggesting a potential selective pressure towards the use of VH5 genes in the repertoire against PIV-3 F, as has been implicated for a class of HIV-1 antibodies [48]. A holistic view of the molecular features underlying antibody reactivity have yet to be described; however, with continued technological advancements in B cell receptor sequencing and the rise of *in silico* antibody engineering [49–51], antigen specificity prediction from antibody sequence alone is expected over time. The data presented here serve to enhance the space of antigen-annotated antibody sequence information, which is a current and major bottleneck for *in silico* modeling and prediction, as discovery efforts to date have been limited to individual pathogen targets, resulting in information for a handful of select antigens.

The results described here provide a better understanding of the BCR sequence features constituting antiviral immunity in the context of twenty antigens. Our analysis of public antibody clonotypes offers novel insights on canonical genetic features of population-level responses and may be used towards the rapid detection of antibodies of a shared reactivity profile from human samples. In addition, antibodies here may serve useful in future investigations as diagnostic tools or potential therapeutics with appropriate functional characterization.

## ACKNOWLEDGMENTS

We thank all members of the Georgiev lab for their support and feedback. We thank David Flaherty, Christian Warren, Olivia Murfield, Emma McLaughlin, and Brittany Matlock from the VUMC Flow Cytometry Shared Resource, for their help with cell sorting. The VUMC Flow Cytometry Shared Resource is supported by the Vanderbilt Ingram Cancer Center (P30 CA68485) and the Vanderbilt Digestive Disease Research Center (DK058404). We thank Angela Jones, Jamie Roberson and Latha Raju with the Vanderbilt Technologies for Advanced Genomics Core (VANTAGE) for providing technical assistance with library production and sequencing. VANTAGE is supported in part by CTSA (5UL1 RR024975-03), the Vanderbilt-Ingram Cancer Center (P30 CA68485), the Vanderbilt Vision Center (P30 EY08126), and NIH/NCRR (G20 RR030956). We also thank the Vanderbilt Institute for Clinical and Translational Research (VICTR, VR54735 to A.A.A.), funded by the National Center for Advancing Translational Science (NCATS) Clinical Translational Science Award (CTSA) Program (5UL1TR002243-03). This work was conducted, in part, using the resources of the Advanced Computing Center for Research and Education (ACCRE) at Vanderbilt University. For work described in this manuscript, I.S.G., A.A.A., M.J.V., P.T.W., and Y.P.S. were supported in part, by the G. Harold and Leila Y. Mathers Charitable Foundation (MF-2107-01851) and NIH R01AI175245 (to I.S.G.). S.A.R and J.S.M were supported in part by Welch Foundation (F-0003-19620604); A.A.A. was supported in part by NIH grant T32 (5T32AI112541-07); P.T.W. was supported in part by NIEHS grant T15 (T15LM007450-19). The funders had no role in the conceptualization or execution of any studies or drafting of the manuscript.

## AUTHOR CONTRIBUTIONS

Conceptualization and Methodology: A.A.A. and I.S.G.; Investigation: A.A.A., M.J.V., P.T.W., Y.P.S., S.A.R., R.A.G., R.S.S. M.C; Writing – Original Draft: A.A.A. and I.S.G.; Writing – Review & Editing: All authors; Funding Acquisition: A.A.A., J.S.M, and I.S.G. Resources: I.S.G; Supervision: A.A.A. and I.S.G.

## METHOD DETAILS

### Healthy donor peripheral blood mononuclear cell (PBMCs) samples

PBMCs donor samples were procured from StemCell Technologies catalog number 70025. Samples arrive frozen and were stored at -135°C until use. Donors are screened for HIV-1, HIV-2, hepatitis B, and hepatitis C within 90 days of collection.

### Cell lines

Ramos B cell lines were engineered from a clone of Ramos Burkitt’s lymphoma that do not display endogenous antibody, and they ectopically express specific surface IgM B cell receptor sequences. The B cell line used here expresses B cell receptor sequences for HIV-specific antibody VRC01. The cells are cultured at 37°C with 5% CO_2_ saturation in complete RPMI, made up of RPMI supplemented with 15% fetal bovine serum, 1% L-Glutamine, and 1% Penicillin/Streptomycin.

### Antigen expression and purification

For the different LIBRA-seq experiments, a total of twenty-one proteins were expressed as recombinant soluble antigens.

Influenza, parainfluenza, HIV-1, and coronavirus antigens were expressed in Expi293F cells by transient transfection in FreeStyle F17 expression media (Thermo Fisher) supplemented to a final concentration of 0.1% Pluronic Acid F-68 and 20% 4 mM L-glutamine using Expifectamine transfection reagent (Thermo Fisher Scientific) cultured for 4-7 days at 8% CO_2_ saturation and 37°C with shaking. After transfection, cultures were centrifuged at 6000 rpm for 20 minutes. Supernatant was filtered with Nalgene Rapid Flow Disposable Filter Units with PES membrane (0.45 or 0.22 μm), and then run slowly over an affinity column at 4C.

Previously described hMPV F A1(NL/1/00) and B2(TN99-419) antigens were expressed in FreeStyle 293-F cells by transient transfection in FreeStyle 293 expression media (Thermo Fisher). Cells were co-transfected at a 4:1 ratio of plasmids encoding human metapneumovirus F and furin, respectively, using polyethylenimine (PEI). Three hours post-transfection, media was supplemented to a final concentration of 0.1% (v/v) Pluronic Acid F-68. After culturing for 6 days at 37°C and 8% CO_2_ saturation, filtered supernatant was concentrated and buffer exchanged to PBS using tangential flow filtration. Samples were then run over a gravity-flow affinity column at room temperature. Previously described RSV F (DS-Cav1) A2 and B9320 antigens were expressed similarly but did not include the Pluronic F-68 supplementation step.

Single chain HIV-1 gp140 SOSIP variant strain BG505 [52] was purified over agarose bound Galanthus nivalis lectin (Vector Laboratories cat no. AL-1243-5). Column was washed with PBS, and protein was eluted with 30 mL of 1 M methyl-a-D-mannopyranoside. The protein elution was buffer exchanged 3X into PBS and concentrated using 30kDa Amicon Ultra centrifugal filter units. Concentrated protein was run on a Superose 6 Increase 10/300 GL on the AKTA FPLC system. Peaks corresponding to trimeric species were identified based on elution volume and SDS-PAGE of elution fractions. Fractions containing pure HIV-1 BG505 Env were pooled.

Recombinant hemagglutinin (HA) proteins from strains all contained the HA ectodomain with a point mutation at the sialic acid-binding site (Y98F), a T4 fibritin foldon trimerization domain, and a hexahistidine-tag. HAs were purified by metal affinity chromatography. 50 mM Tris pH 8.0 and 350 mM NaCl was added to the clarified supernatant using concentrated solutions of 1 M Tris pH 8.0 and 5 M NaCl, respectively. For each liter of supernatant, 4 ml of Ni Sepharose Excel resin (GE) bed was washed with PBS using a gravity column and then added to the buffered supernatant, followed by overnight rocking at 4°C. The resin was collected 16–24 h later using a gravity column, then washed once with 50 mM Tris pH 8.0, 500 mM NaCl, 5 mM imidazole prior to elution of His-tagged protein using 50 mM Tris pH 8.0, 500 mM NaCl, 300 mM imidazole. Eluates were concentrated and applied to a HiLoad 16/600 Superdex 200 pg column or a Superdex 200 Increase 10/300 GL column pre-equilibrated with PBS for preparative size exclusion chromatography. Peaks corresponding to trimeric species were identified based on elution volume and SDS-PAGE of elution fractions. Fractions containing pure HA were pooled.

Parainfluenza virus type 3 prefusion stabilized F ectodomain (PDB: 6MJZ) was purified by nickel affinity chromatography using an equilibrated, 1 mL pre-packed HisTrap HP column (GE Healthcare, IL, USA). The column was washed with 15 mL of binding buffer (20 mM sodium phosphate, 0.5 M NaCl, 0.3 M imidazole, pH 7.4), and purified protein was eluted from the column with 15 mL of binding buffer supplemented with 0.5 M Imidazole. Protein was concentrated, buffer exchanged into PBS, and run on a Superose 6 Increase 10/300 GL on the AKTA FPLC system. Peaks corresponding to trimeric species were identified based on elution volume and SDS-PAGE of elution fractions. Fractions containing pure PIV-3 F were pooled.

SARS-CoV-2 S Hexapro Wuhan strain, HCoV-OC43 S, and HCoV-HKU1-S-2P were purified over equilibrated, 1 mL pre-packed StrepTrap XT column (Cytiva Life Sciences). The column was washed with 15 mL of binding buffer (100 mM Tris-HCl, 150 mM NaCl, 1 mM EDTA, pH 8.0), and purified protein was eluted from the column with 10 mL of binding buffer supplemented with 2.5 mM desthiobiotin. Protein was concentrated, buffer exchanged into PBS, and run on a Superose 6 Increase 10/300 GL on the AKTA FPLC system. Peaks corresponding to trimeric species were identified based on elution volume and SDS-PAGE of elution fractions. Fractions containing pure spike were pooled.

Stabilized ectodomains of hMPV F subtypes A1 and B2 ([38], as well as RSV strains A2 ([53, 54]and B9320 F (DS-Cav1)[55], were purified over Strep-Tactin Sepharose resin (IBA Lifesciences) in a gravity column. The resin was washed with 4 column volumes of PBS, and purified protein was eluted from the column with 4 column volumes of 100 mM Tris pH 8.0, 150 mM NaCl, 1 mM EDTA, 2.5 mM desthiobiotin. Eluate was concentrated and run on a Superose 6 increase 10/300 GL column pre-equilibrated with PBS (except for RSV A2 F which was pre-equilibrated with 2 mM Tris pH 8.0, 200 mM NaCl, and 0.02% NaN_3_) for preparative size exclusion chromatography. Peaks corresponding to trimeric species were identified based on elution volume and SDS-PAGE of elution fractions. Fractions containing pure fusion protein were pooled.

DENV-1, DENV-2, DENV-3, DENV-4 E glycoproteins were recombinantly expressed and purified as previously described [56].

All proteins were quantified using UV/vis spectroscopy. Antigenicity of proteins was characterized by ELISA with known monoclonal antibodies specific for that antigen. Proteins were frozen and stored at -80°C until use.

### Biotinylation of antigens

Protein antigens were biotinylated using EZ-link Sulfo-NHS-Biotin No-Weigh kit (Thermo Fisher) according to manufacturer’s instructions. A 50:1 biotin-to-protein molar ratio was used for all reactions.

### Oligonucleotide barcodes

We used oligos that possess a 15 bp antigen barcode, a sequence capable of annealing to the template switch oligo that is part of the 10X bead-delivered oligos and contain truncated TruSeq small RNA read 1 sequences in the following structure: 5′- CCTTGGCACCCGAGAATTCCANNNNNNNNNNNNNNNCCCATATAAGA*A*A-3’, where Ns represent the antigen barcode. Oligos were ordered from Sigma-Aldrich and IDT with a 5’ amino modification and HPLC purified. The following antigen barcodes were used: CCGTCCTGATAGATG (HKU1), GTGTGTTGTCCTATG (OC43), TCACAGTTCCTTGGA (SARS-CoV-2), ACAATTTGTCTGCGA (RSV A), CAGGTCCCTTATTTC (RSV B), CAGCCCACTGCAATA (hMPV A), AACCTTCCGTCTAAG (hMPV B), CAGATGATCCACCAT (PIV-3), TACGCCTATAACTTG (H1 NC/99), TGGTAACGACAGTCC (H1 MI/15), AATCACGGTCCTTGT (H3 Perth), TCATTTCCTCCGATT (H3 HK/68), TCCTTTCCTGATAGG (H5 VN/04), CAGTAAGTTCGGGAC (H7 AN/13), ATTCGCCTTACGCAA (H9 HK/09), ATCGTCGAGAGCTAG (H10 JD/13), TGTGTATTCCCTTGT (DENV 1), CTTCACTCTGTCAGG (DENV 2), CAGTAGATGGAGCAT (DENV 3), GGTAGCCCTAGAGTA (DENV 4) and TAACTCAGGGCCTAT (HIV-1)

### Conjugation of oligonucleotide barcodes to antigens

For each antigen, a unique DNA barcode was directly conjugated to the antigen using a SoluLINK Protein-Oligonucleotide Conjugation kit (TriLink, S-9011) according to manufacturer’s protocol. Briefly, the oligo and protein were desalted, and then the amino-oligo was modified with the 4FB crosslinker, and the biotinylated antigen protein was modified with S-HyNic. Then, the 4FB-oligo and the HyNic-antigen were mixed together. This causes a stable bond to form between the protein and the oligonucleotide. The concentration of the antigen-oligo conjugates was determined by a BCA assay, and the HyNic molar substitution ratio of the antigen-oligo conjugates was analyzed using the NanoDrop according to the Solulink protocol guidelines. AKTA FPLC was used to remove excess oligonucleotide from the protein-oligo conjugates, which were also checked using SDS-PAGE with a silver stain.

### Enrichment of antigen-specific B cells

For a given sample, cells were mixed ∼4% (of total viable PBMCs) with VRC01 Ramos cells, to serve as an internal negative control. Cell mixtures were stained and mixed with fluorescently labeled DNA-barcoded antigens and other antibodies, and then sorted using fluorescence activated cell sorting (FACS). First, cells were counted, and viability was assessed using Trypan Blue. Then, cells were washed with DPBS supplemented with 1% Bovine serum albumin (BSA) through centrifugation at 300 g for 5 minutes. Cells were resuspended in PBS-BSA and stained with a variety of cell markers. These markers included Ghost Red 780, CD14-APCCy7, CD3-FITC, CD19-BV711, and IgG-PECy5. Additionally, antigen-oligo conjugates were added to the stain. After a 30-minute incubation in the dark on ice, the cells were washed again three times with DPBS-BSA at 300 g for 5 minutes. Then, the cells were incubated for 15 minutes in the dark on ice with Streptavidin-PE and Streptavadidin-BV421 to label cells with bound antigen. Cells were then resuspended in DPBS and sorted on the cell sorter. Antigen positive cells were bulk sorted and then delivered to the Vanderbilt VANTAGE sequencing core at an appropriate target concentration for 10X Genomics library preparation and subsequent sequencing. FACS data were analyzed using FlowJo.

### 10X Genomics single cell processing and next generation sequencing

Single-cell suspensions were loaded onto the Chromium Controller microfluidics device (10X Genomics) and processed using the B cell Single Cell V(D)J solution according to manufacturer’s suggestions for a target capture of 10,000 B cells per 1/8 10X cassette for B cells. Slight modifications were made to intercept, amplify, and purify the antigen barcode libraries, as previously described [24].

### Sequence processing and bioinformatics analysis

We followed our established pipeline, which takes paired-end FASTQ files of oligonucleotide libraries as input, to process and annotate reads for cell barcodes, unique molecular identifiers (UMIs) and antigen barcodes, resulting in a cell barcode-antigen barcode UMI count matrix [24]. B cell receptor contigs were processed using CellRanger 3.1.0 (10x Genomics) and GRCh38 Human V(D)J 7.0.0 as reference, while the antigen barcode libraries were also processed using CellRanger (10x Genomics). The cell barcodes that overlapped between the two libraries formed the basis of the subsequent analysis. Cell barcodes that had only non-functional heavy chain sequences as well as cells with multiple functional heavy chain sequences and/or multiple functional light chain sequences, were eliminated, reasoning that these may be multiplets. We also aligned the B cell receptor contigs (filtered_contigs.fasta file output by CellRanger, 10x Genomics) to IMGT reference genes using HighV-Quest [57]. The output of HighV-Quest was parsed using ChangeO [58] and combined with an antigen barcode UMI count matrix. Finally, we determined the LIBRA-seq score for each antigen in the library for every cell as previously described [24].

### Filtering Pipeline

To adjust for noise due to ambient antigen barcode capture and non-specific binding interactions, a negative binomial mixture model was developed to calculate the probability of a given UMI count being considered signal or noise. This approach leveraged the VRC01-BG505 negative control system to fit a mixed model for the distribution of UMI counts for each antigen within each individual sample. For each antigen, the distribution of UMI counts for the antigen bound to VRC01 was assumed to be technical noise. VRC01 cells were selected using a 95% identity threshold to the VRC01 CDRH3 nucleotide sequence using Levenshtein distance. A mixed binomial distribution then was fit to the distribution of UMIs representing binding to the donor (non-VRC01) cells for that antigen. After fitting, probability mass functions (PMF) were then generated from the optimized parameters, and Baye’s theorem was used to calculate the probability of each UMI count being in the ‘signal’ component of the mixed distribution. This is an adaptation of a mixed gaussian model, using the negative binomial distributions to cluster the UMI counts into each of the two distributions. Finally, the probability of being signal was used for filtering, with a 90% probability threshold included in addition to the LIBRA-seq score (LSS) threshold of 1.VRC01 cells were not recovered from Donor 3 (8365-3) and this sample therefore did not utilize this mixed modeling filtering approach.

After generating probabilities of UMI counts being signal in the raw counts using the mixed model described above, the following filtering steps were implemented to further filter the cells using a custom Python script. VRC01 cells were removed, along with cells that bound to the BG505 negative control antigen (LSS ≥1). To account for technical artifacts resulting from LSS calculated for antigens with very low counts across all cells, UMI counts that were less than 10 but yielded LSS ≥ 1 were set to the minimum LSS for that antigen. For defining binding families and type-specificities, binding was defined as having a LSS≥1. Cells that bound to more than one viral family were considered polyreactive and removed, along with cells with LSS <1 for all antigens in the experiment. Only IgG isotype cells were retained for the systematic repertoire (non-public) analyses.

Following in-vitro validation by ELISA, filtering thresholds were modified to inform the systems and clonotype analysis. For specific antibodies tested by ELISA, LSS for these cells were updated to reflect the assay results. A further UMI threshold for Dengue antigens (DENV E 3 and DENV E 4) was also implemented due to the high false positive LSS rate observed, with UMI counts less than 400 for DENV E 3 and less than 120 for DENV E 4 considered non-binding.

### V(D)J Sequence Analysis

After pre-processing and filtering, the V(D)J sequences and annotations were further analyzed using custom scripts written in Python 3.10, utilizing NumPy 1.23 and Pandas 1.4.3 for data wrangling, Matplotlib 3.6.2 and Seaborn 0.12.2 for visualization, and Scipy 1.8.1.

### Public antibody analysis

The reference library combined paired heavy chain-light chain human BCR sequences from published sources [41, 42]. We used custom Python scripts utilizing NumPy and Pandas to determine public clones between our data and the reference library. Public clones were identified on the basis of matching CDRH3 and CDRL3 amino acid lengths, a minimum of 70% amino-acid sequence identity in both CDRH3 and CDRL3 regions and matching heavy and light variable (V) and joining (J) gene usage. Levenshtein distance was used for comparison in CDR3 identities. Shared clusters were then assigned if sequences originating from a minimum of two donors shared viral antigen specificity, maintained a minimum of 70% amino acid identity in both CDR regions, and leveraged the same pair of V and J genes.

### High throughput antibody micro expression

Microscale transfection was performed with Expi293F cells in FreeStyle F17 expression media (Thermo Fisher) supplemented to a final concentration of 0.1% Pluronic Acid F-68 and 20% 4 mM L-glutamine using Expifectamine transfection reagent (Thermo Fisher Scientific). Briefly, synthesized antibody-encoding DNA (1.6 μg per transfection) was added to OptiMem, incubated with ExpiFectamine transfection reagent, and added to 800 µl of Expi293 cell cultures in deep 96-well blocks. The plates were incubated on an orbital shaker at 900 r.p.m. with an orbital diameter of 3 mm at 37 °C in 8% CO2. Culture supernatants were collected at day 5 post transfection. Blocks were centrifuged at max speed and supernatant was carefully removed and immediately used for ELISA.

### Enzyme linked immunosorbent assay (ELISA)

Soluble protein was plated at 2 ug/ml overnight at 4°C. The next day, plates were washed three times with PBS supplemented with 0.05% Tween20 (PBS-T) and coated with 1% BSA in PBS-T. Plates were incubated for one hour at room temperature and then washed three times with PBS-T. Supernatant from antibody microscale transfection was then plated and incubated at room temperature for one hour and then washed three times in PBS-T. The secondary antibody, goat anti-human IgG conjugated to peroxidase, was added at 1:10,000 dilution in 1% BSA in PBS-T to the plates, which were incubated for one hour at room temperature. Plates were washed three times with PBS-T and then developed by adding TMB substrate to each well. The plates were incubated at room temperature for five minutes, and then 1 N sulfuric acid was added to stop the reaction. Plates were read at 450 nm. ELISAs were repeated 2 or more times.

---

**Supplemental Figure 1:**
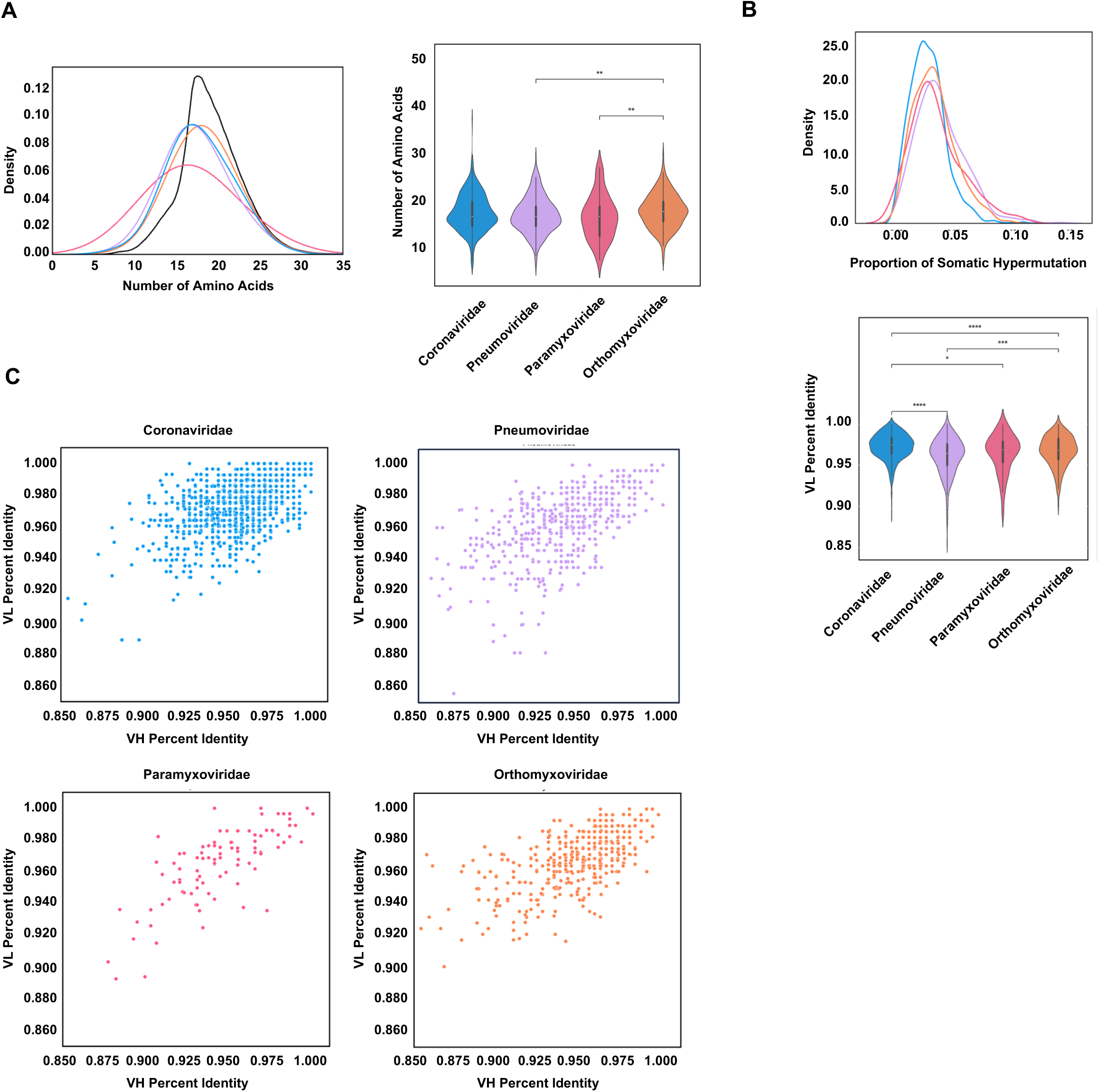
CDRH3 length and somatic hypermutation of viral antigen specific B cells. 1A: CDRH3 length represented as the frequency within each antigen specificity category. Previously published, unselected repertories are shown in black. Violin plot width is proportional to the fraction of B cells with the indicated CDRH3 length. Two sided Mann-Whitney-Wilcoxon test with Bonferroni correction used to calculate significance. Coronaviridae vs. Pneumoviridae: p= 1.038^-01^, Pneumoviridae vs. Paramyxoviridae: p= 8.767^-01^, Paramyxoviridae vs. Orthomyxoviridae: p= 4.052^-03^, Coronaviridae vs. Paramyxoviridae: p= 5.075^-02^, Pneumoviridae vs. Orthomyxoviridae: p= 1.292^-03^, Coronaviridae vs. Orthomyxoviridae: p= 3.050^-0^. 1B: Frequency of light chain variable (VL) somatic hypermutation represented as 1-VL identity calculated at the amino acid level. Violin plot width is proportional to the fraction of B cells with the indicated proportion of VL somatic hypermutations. Two sided Mann-Whitney-Wilcoxon test with Bonferroni correction used to calculate significance. Coronaviridae vs. Pneumoviridae: p= 1.118^-23^, Pneumoviridae vs. Paramyxoviridae: p= 2.870^-01^, Paramyxoviridae vs. Orthomyxoviridae: p= 1.000, Coronaviridae vs. Paramyxoviridae: p= 1.636^-02^, Pneumoviridae vs. Orthomyxoviridae: p= 4.598^-04^, Coronaviridae vs. Orthomyxoviridae: p= 2.601^-06^. 1C: Correlation between the proportion of heavy chain variable gene somatic hypermutation (x axis) and light chain variable gene somatic hypermutation (y axis) of B cells for each antigen specificity category. Coronaviridae spearman correlation coefficient 0.47 p = 7.2^-63^. Pneumoviridae spearman correlation coefficient 0.57 p= 7.9^-48^. Paramyxoviridae spearman correlation 0.74 p= 2.1^-21^. Orthomyxoviridae spearman correlation coefficient 0.62 p=8.7^-47^.

**Supplemental Figure 2:**
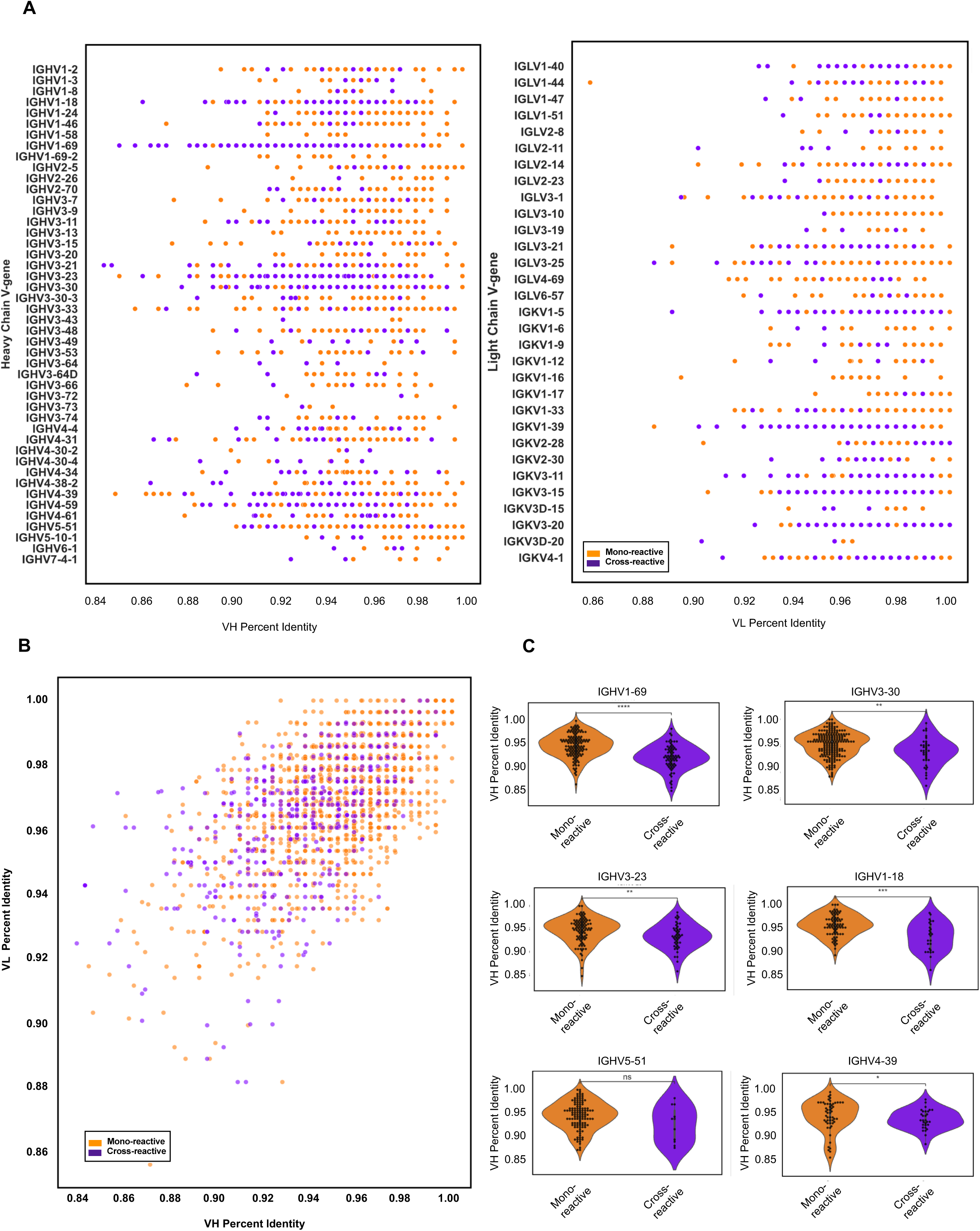
Variable gene somatic hypermutation of mono-reactive and cross-reactive B cells. 2A: Dot plot indicating VH percent identity for IGHV genes leveraged by mono-reactive and cross-reactive cells. Dot plot indicating VL percent identity for IGL(K)V genes leveraged by mono-reactive and cross-reactive cells. 2B: Correlation between the VH percent identity (x axis) and VL percent identity (y axis). Mono-reactive spearman correlation coefficient 0.57, p= 1.4^-152^. Cross-reactive spearman correlation coefficient 0.54, p=1.2^-36^. 2C: VH percent identity for the most represented IGHV genes leveraged by both mono-reactive and cross-reactive cells. In order of most represented to least: IGHV1-69, IGHV3-30, IGHV3-23, IGHV1-18, IGHV5-51, IGHV4-39. IGHV1-69 p = 1.584^-10^, IGHV3-30 p = 1.388^-03^, IGHV3-23 p = 4.028^-03^, IGHV1-18 p = 3.367^-04^, IGHV5-51 p = 1.319^-01^, IGHV4-39 = 4.476^-02^

**Supplemental Figure 3:**
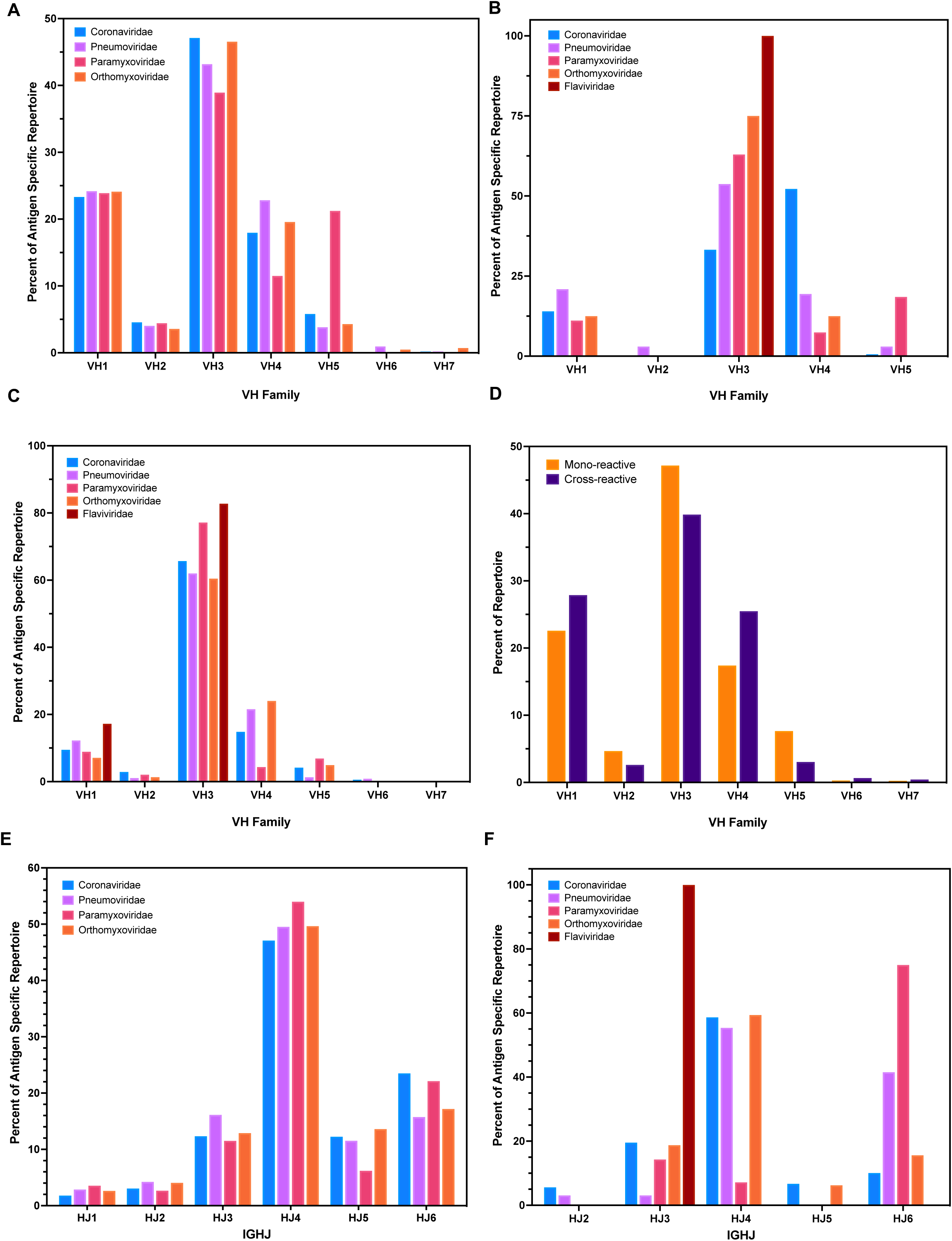
Variable (V) and joining (J) gene segment usage frequencies. 3A: Observed frequencies of V gene families from the IgG restricted dataset, represented as frequency of segment use within each antigen specificity category. 3B: Observed frequencies of V gene families from identified public clonotypes matching on heavy chain and light chain similarity criteria, represented as frequency of segment use within each antigen specificity category. 3C:Observed frequencies of V gene families from identified public clonotypes matching on heavy chain similarity criterion, represented as frequency of segment use within each antigen specificity category. 3D:Observed frequencies of V gene families from the IgG restricted dataset, partitioned by reactivity pattern, represented as frequency of segment use within each reactivity category. 3E:Observed frequencies of J genes from the IgG restricted dataset, partitioned by reactivity pattern, represented as frequency of segment use within each reactivity category. 3F:Observed frequencies of J genes from identified public clonotypes matching on heavy chain and light chain similarity criteria, represented as frequency of segment use within each reactivity category.

**Supplemental Figure 4:**
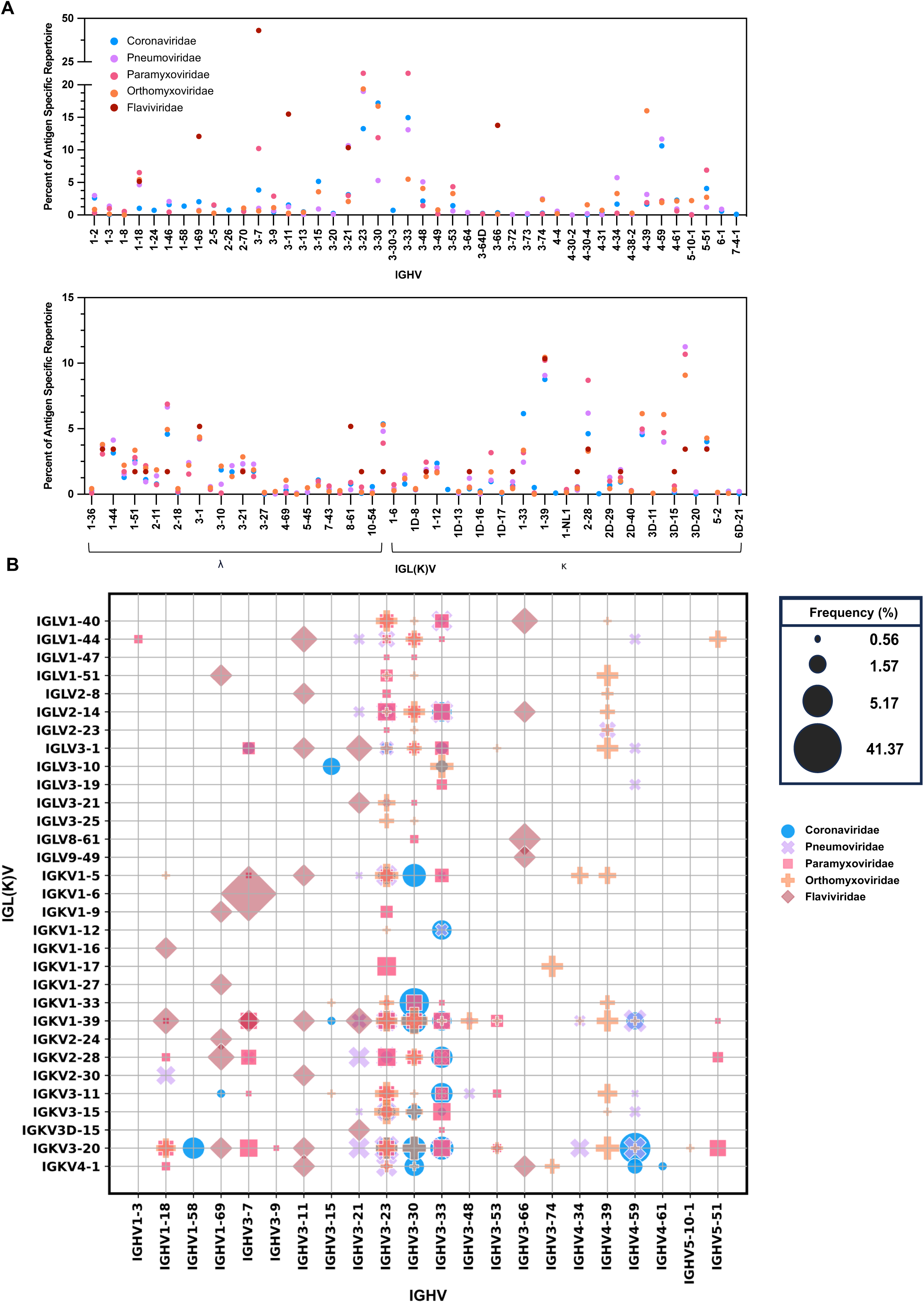
B cell characteristics of viral antigen public clonotypes identified by heavy chain similarity criteria. 4A: IGHV and IGL(K)V gene usage in public clonotypes specific to Coronaviridae (blue), Pneumoviridae (purple), Paramyxoviridae (pink), Orthomyxoviridae (orange), and Flaviviridae (red) antigens. Gene usage is represented as frequency of gene use within antigen specificity category. Circles are superimposed. Zeros are not plotted. 4B: The frequency of different IGHV:IGL(K)V gene pairs for public clonotypes specific to Coronaviridae (blue circle), Pneumoviridae (purple cross), Paramyxoviridae (pink square), Orthomyxoviridae (orange plus), Flaviviridae (red diamond) antigens. The size of each data point represents the frequency of the corresponding IGHV:IGK(L)V pair within its antigen specificity category. Pairs above 0.5% are represented.

**Supplemental Table 1:**
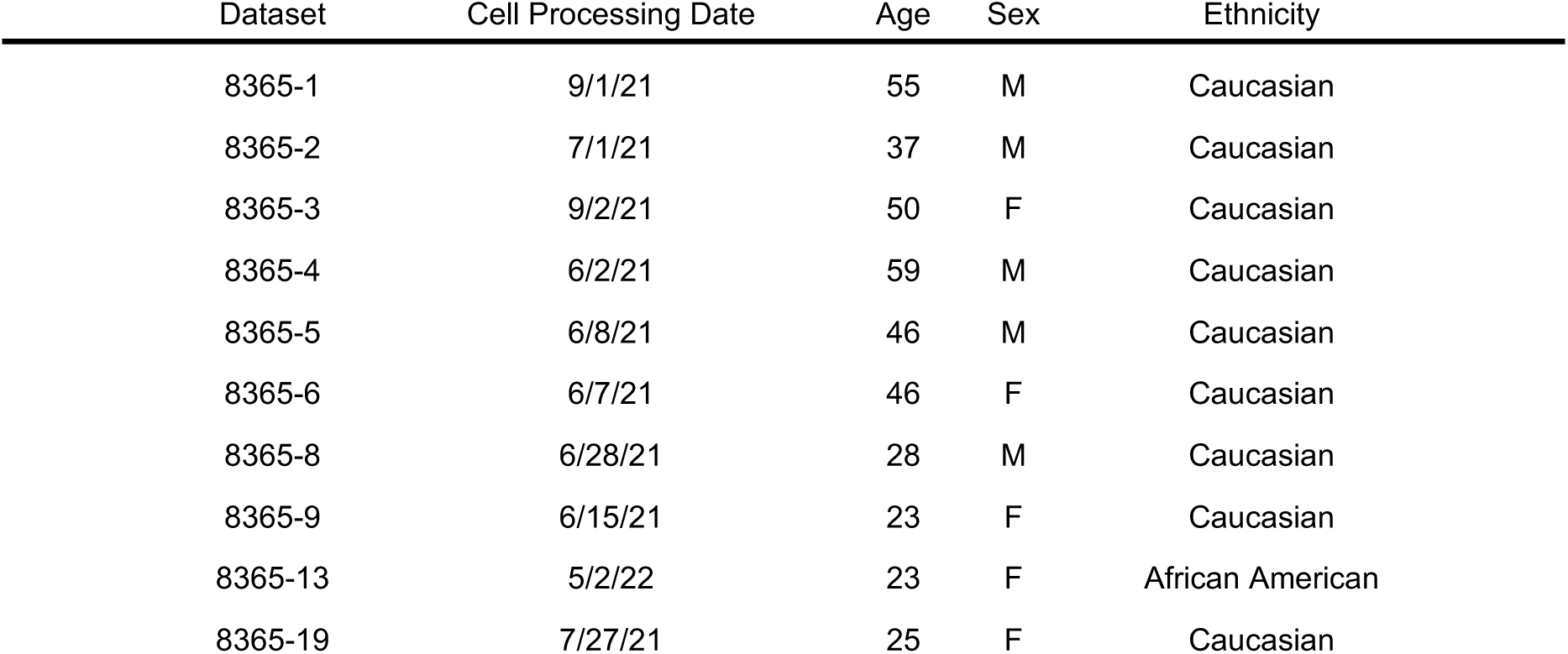
Donor peripheral mononuclear blood cell (PBMC) data. PBMC samples were procured from StemCell Technologies. Donor cell processing date, age, sex, and ethnicity are listed for each donor.

| Dataset | Cell Processing Date | Age | Sex | Ethnicity |
| --- | --- | --- | --- | --- |
| 8365-1 | 9/1/21 | 55 | M | Caucasian |
| 8365-2 | 7/1/21 | 37 | M | Caucasian |
| 8365-3 | 9/2/21 | 50 | F | Caucasian |
| 8365-4 | 6/2/21 | 59 | M | Caucasian |
| 8365-5 | 6/8/21 | 46 | M | Caucasian |
| 8365-6 | 6/7/21 | 46 | F | Caucasian |
| 8365-8 | 6/28/21 | 28 | M | Caucasian |
| 8365-9 | 6/15/21 | 23 | F | Caucasian |
| 8365-13 | 5/2/22 | 23 | F | African American |
| 8365-19 | 7/27/21 | 25 | F | Caucasian |

**Supplemental Table 2:** Identified public clonotypes. Number of sequences identified as public from the reference library and the LIBRA-seq datasets are shown for each antigen specificity category.

| HC+LC Elements | Coronaviridae | Pneumoviridae | Paramyxoviridae | Orthomyxoviridae | Flaviviridae |
| --- | --- | --- | --- | --- | --- |
| Reference library | 290 | 47 | 16 | 20 | 14 |
| LIBRA-seq dataset | 68 | 20 | 11 | 12 | 2 |
| HC Elements |  |  |  |  |  |
| Reference library | 4583 | 3019 | 1026 | 1263 | 50 |
| LIBRA-seq dataset | 416 | 244 | 78 | 135 | 8 |

**Supplemental Table 3:** Public cluster specificity and isotype. Public clusters identified on basis of matching CDRH3 and CDRL3 length, a minimum of 70% amino acid identity in CDRH3 and CDRL3, and the same pair of V and J gene use in heavy and light chains. Number of sequences and donors within each cluster are represented. Antigen specificity and isotype determination afforded by LIBRA-seq and next generation sequencing.

| Cluster Number | Sequences | Donors | Viral Family | Antigen Specificity | Isotype | Cluster Number | Sequences | Donors | Viral Family | Antigen Specificity | Isotype |
| --- | --- | --- | --- | --- | --- | --- | --- | --- | --- | --- | --- |
| 1 | 2 | 2 | Coronaviridae | SARS-CoV-2 spike | IgG3 | 48 | 5 | 3 | Coronaviridae | SARS-CoV-2 spike | IgG1 |
| 2 | 4 | 3 | Paramyxoviridae | PIV-3 F | IgG1 | 49 | 2 | 2 | Paramyxoviridae | PIV-3 F | IgM |
| 3 | 11 | 4 | Pneumoviridae | RSV A F, RSV B F | IgG2 | 50 | 4 | 2 | Coronaviridae | SARS-CoV-2 spike | IgM |
| 4 | 2 | 2 | Orthomyxoviridae | H3 Perth HA | IgG1 | 51 | 5 | 2 | Pneumoviridae | hMPV B F | IgM |
| 5 | 3 | 2 | Pneumoviridae | MPV A F | IgG1 | 52 | 2 | 2 | Coronaviridae | SARS-CoV-2 spike | IgG1 |
| 6 | 2 | 2 | Coronaviridae | SARS-CoV-2 spike | IgG1 | 53 | 2 | 2 | Pneumoviridae | RSV A F | IgM |
| 7 | 2 | 2 | Coronaviridae | SARS-CoV-2 spike | IgG1 | 54 | 7 | 3 | Orthomyxoviridae | H7 AN13 HA | IgG1 |
| 8 | 3 | 2 | Coronaviridae | SARS-CoV-2 spike | IgA1 | 55 | 2 | 2 | Pneumoviridae | RSV A F, RSV B F | IgM |
| 9 | 30 | 3 | Coronaviridae | SARS-CoV-2 spike | IgG1 | 56 | 2 | 2 | Pneumoviridae | hMPV A F, hMPV B F | IgG2 |
| 10 | 2 | 2 | Orthomyxoviridae | H1 NC99 HA | IgG1 | 57 | 2 | 2 | Coronaviridae | SARS-CoV-2 spike | IgG1 |
| 11 | 2 | 2 | Coronaviridae | SARS-CoV-2 spike | IgG1 | 58 | 5 | 3 | Paramyxoviridae | PIV-3 F | IgA1 |
| 12 | 3 | 2 | Coronaviridae | SARS-CoV-2 spike | IgG1 | 59 | 2 | 2 | Coronaviridae | SARS-CoV-2 spike | IgG1 |
| 13 | 4 | 2 | Coronaviridae | SARS-CoV-2 spike | IgG1 | 60 | 2 | 2 | Paramyxoviridae | PIV-3 F | IgG2 |
| 14 | 2 | 2 | Pneumoviridae | RSV A F | IgG1 | 61 | 14 | 2 | Flaviviridae | DENV 4 E | n/a |
| 15 | 2 | 2 | Flaviviridae | Dengue E 1 | IgM | 62 | 13 | 5 | Coronaviridae | OC43 spike | IgM |
| 16 | 2 | 2 | Coronaviridae | SARS-CoV-2 spike | IgM | 63 | 2 | 2 | Coronaviridae | SARS-CoV-2 spike | IgG1 |
| 17 | 2 | 2 | Orthomyxoviridae | H1 MI15 HA | IgG1 | 64 | 2 | 2 | Coronaviridae | SARS-CoV-2 spike | IgG1 |
| 18 | 3 | 2 | Coronaviridae | SARS-CoV-2 spike | IgG1 | 65 | 2 | 2 | Pneumoviridae | RSV B F | IgG2 |
| 19 | 2 | 2 | Orthomyxoviridae | H1 MI15 HA | IgG1 | 66 | 2 | 2 | Pneumoviridae | RSV A F | IgM |
| 20 | 2 | 2 | Coronaviridae | SARS-CoV-2 spike | IgM | 67 | 2 | 2 | Orthomyxoviridae | H3 HK68 HA | IgG1 |
| 21 | 8 | 3 | Coronaviridae | SARS-CoV-2 spike | IgG1 | 68 | 2 | 2 | Paramyxoviridae | PIV-3 F | IgM |
| 22 | 8 | 5 | Coronaviridae | SARS-CoV-2 spike | IgG1 | 69 | 2 | 2 | Orthomyxoviridae | H1 NC99 HA | IgM |
| 23 | 2 | 2 | Coronaviridae | SARS-CoV-2 spike | IgG1 | 70 | 4 | 3 | Pneumoviridae | RSV A F | IgM |
| 24 | 3 | 2 | Coronaviridae | SARS-CoV-2 spike | IgG4 | 71 | 2 | 2 | Orthomyxoviridae | H3 HK68 HA | IgG1 |
| 25 | 3 | 3 | Coronaviridae | SARS-CoV-2 spike | IgG1 | 72 | 10 | 2 | Coronaviridae | SARS-CoV-2 spike | IgG1 |
| 26 | 4 | 2 | Pneumoviridae | RSV B F | IgM | 73 | 2 | 2 | Paramyxoviridae | PIV-3 F | IgM |
| 27 | 4 | 2 | Coronaviridae | SARS-CoV-2 spike | IgG1 | 74 | 2 | 2 | Pneumoviridae | hMPV A F, hMPV B F | IgG1 |
| 28 | 2 | 2 | Coronaviridae | SARS-CoV-2 spike | IgG1 | 75 | 2 | 2 | Coronaviridae | SARS-CoV-2 spike | IgG1 |
| 29 | 8 | 3 | Coronaviridae | SARS-CoV-2 spike | IgG1 | 76 | 2 | 2 | Coronaviridae | SARS-CoV-2 spike | IgM |
| 30 | 5 | 2 | Pneumoviridae | RSV A F | IgM | 77 | 5 | 3 | Pneumoviridae | hMPV A F | IgM |
| 31 | 2 | 2 | Orthomyxoviridae | H1 MI15 HA | IgG1 | 78 | 5 | 4 | Coronaviridae | SARS-CoV-2 spike | IgG2 |
| 32 | 2 | 2 | Pneumoviridae | RSV A F | IgM | 79 | 3 | 3 | Pneumoviridae | RSV A F, RSV B F | IgM |
| 33 | 2 | 2 | Paramyxoviridae | PIV-3 F | IgM | 80 | 18 | 3 | Coronaviridae | SARS-CoV-2 spike | IgG1 |
| 34 | 2 | 2 | Coronaviridae | SARS-CoV-2 spike | IgG1 | 81 | 93 | 6 | Coronaviridae | SARS-CoV-2 spike | IgG1 |
| 35 | 4 | 2 | Coronaviridae | SARS-CoV-2 spike | IgG1 | 82 | 3 | 2 | Coronaviridae | SARS-CoV-2 spike | IgG1 |
| 36 | 3 | 3 | Coronaviridae | SARS-CoV-2 spike | IgG1 | 83 | 2 | 2 | Coronaviridae | SARS-CoV-2 spike | IgG1 |
| 37 | 4 | 2 | Orthomyxoviridae | H1 MI15 HA | IgG1 | 84 | 4 | 2 | Coronaviridae | SARS-CoV-2 spike | IgG1 |
| 38 | 2 | 2 | Pneumoviridae | RSV A F | IgM | 85 | 16 | 3 | Coronaviridae | SARS-CoV-2 spike | IgG1 |
| 39 | 14 | 2 | Coronaviridae | SARS-CoV-2 spike | IgG1 | 86 | 2 | 2 | Coronaviridae | SARS-CoV-2 spike | IgG1 |
| 40 | 2 | 2 | Coronaviridae | SARS-CoV-2 spike | IgG1 | 87 | 2 | 2 | Coronaviridae | SARS-CoV-2 spike | IgM |
| 41 | 2 | 2 | Coronaviridae | SARS-CoV-2 spike | IgG1 | 88 | 2 | 2 | Pneumoviridae | hMPV A F | IgM |
| 42 | 3 | 2 | Coronaviridae | SARS-CoV-2 spike | IgG1 | 89 | 2 | 2 | Paramyxoviridae | PIV-3 F | IgG1 |
| 43 | 2 | 2 | Coronaviridae | SARS-CoV-2 spike | IgG1 | 90 | 4 | 2 | Paramyxoviridae | PIV-3 F | IgG1 |
| 44 | 4 | 2 | Coronaviridae | SARS-CoV-2 spike | IgG1 | 91 | 3 | 2 | Orthomyxoviridae | H3 Perth HA | IgG1 |
| 45 | 2 | 2 | Coronaviridae | SARS-CoV-2 spike | IgG1 | 92 | 3 | 2 | Coronaviridae | SARS-CoV-2 spike | n/a |
| 46 | 4 | 3 | Coronaviridae | SARS-CoV-2 spike | IgG1 | 93 | 2 | 2 | Coronaviridae | SARS-CoV-2 spike | IgG1 |
| 47 | 2 | 2 | Orthomyxoviridae | H1 NC99 HA | IgM | 94 | 2 | 2 | Coronaviridae | SARS-CoV-2 spike | IgG1 |
|  |  |  |  |  |  | 95 | 2 | 2 | Paramyxoviridae | PIV-3 F | IgM |

